# Two mutations in the same MYC-bHLH transcription factor cause segregation of purple coloration of stolons and seed heads in *Zoysia japonica* x *Zoysia matrella* F_2_ and F_1_ populations

**DOI:** 10.1101/2025.08.04.668483

**Authors:** Shreena Pradhan, Jianxin Zhao, John J Spiekerman, Emma M Bennetzen, Sameer Khanal, Xingwang Yu, Susana Milla-Lewis, Joann Conner, Brian M Schwartz, Katrien M Devos

## Abstract

Anthocyanins play diverse roles in plants, including attracting pollinators and protecting cells from oxidative damage. In zoysiagrass, a warm season turfgrass, their accumulation in seed heads and stolons can decrease the aesthetic appeal. In this study, a high-density genetic map with ∼8000 single nucleotide polymorphism (SNP) markers organized into 20 linkage groups was generated in a *Zoysia japonica* acc. Meyer x *Zoysia matrella* acc. PI 231146 F_2_ population. Using this genetic map, a large-effect quantitative trait locus (QTL) for anthocyanin variation in stolons and seed heads was mapped to chromosome 12 (*PP* locus). Variant analysis of a candidate gene for *PP*, *Zjn_sc00004.1.g07010.1.sm.mk*, which encodes a MYC-bHLH transcription factor that regulates anthocyanin biosynthesis, revealed a SNP at an exon-intron boundary in Meyer that led to intron retention. Interestingly, an F_1_ population derived from the same parents segregated for seed head color but uniformly displayed purple stolons. Seed head color in the F_1_ population co-mapped with the *PP* locus which, combined with genotypic and yeast two-hybrid analyses, revealed that a SNP in PI 231146 leading to an Ala163Ser substitution in the MYB-interacting N-terminal domain of the same MYC-bHLH transcription factor was likely causal. The Ala163Ser substitution affected interaction of MYC-bHLH with MYB in a MYB-dependent manner. The identified mutations can be exploited to develop cultivars with green seed heads and stolons. The high-marker-density interspecific *Z. japonica* x *Z. matrella* F_2_ genetic map also provides a robust tool for identifying genomic regions and genes of agronomic interest that differentiate the two species.

## Introduction

Zoysiagrass is the common name for several introduced perennial sod-forming species in the genus *Zoysia* with the three primary species used as turf being *Z. japonica* Steud., *Z. matrella* (L.) Merr., and *Z. pacifica* (Goudswaard) M. Hotta & S. Kuroki [1]. As a C4 grass, zoysiagrass thrives primarily in warm-humid climatic zones, and is characterized by low fertility requirements and a notable tolerance to various stresses such as drought and salinity [2]. It is extensively utilized across the southern United States on golf courses, home lawns, athletic fields, and other recreation sites. The increasing popularity of zoysiagrass can be attributed to the development of varieties with good shade and wear tolerance, high cold hardiness and high shoot density to suit different environments and uses. [3]. In 2023, the zoysiagrass market in the US was valued at USD 42.6 million, with an anticipated growth rate of 4.3% per annum (https://www.verifiedmarketresearch.com/). Despite its significant commercial value, limited studies in zoysiagrass have focused on the genetic understanding of traits, stress tolerance mechanisms, and marker-based trait improvement strategies.

Zoysiagrass is an allotetraploid (2*n* = 4*x* = 40; C=491 Mb) protogynous turfgrass in which both cross- and self-pollinations are possible. The three primary species can be intercrossed, and hybrid cultivars combining favorable characteristics from the parental species have been released [3]. Several genomic resources are available for zoysiagrass, including a chromosome-level assembly for *Z. japonica* acc. “Nagirizaki” (ZJN_r1.1_pseudomol), and draft assemblies for *Z. matrella* acc. “Wakaba” and *Z. pacifica* acc. “Zanpa” [4]. SNP-based linkage maps generated in *Z. japonica* F_1_ populations [5, 6] as well as in a *Z. matrella* F_1_ population [7] have been documented. However, comprehensive genetic studies, particularly in interspecific F_2_ mapping populations that are suitable for mapping interspecific trait variation, are limited. Transcriptome data are available for *Z. japonica* [8–10] as well as *Z. matrella* [11–14].

Turf quality is a pivotal trait influencing the commercial and aesthetic value of turfgrasses, including zoysiagrass, with uniform color being a critical factor in its appeal. The majority of zoysiagrass cultivars have purple stolons and spikes due to the accumulation of anthocyanins. Anthocyanin, a phenolic compound, imparts vivid colors to attract pollinators and seed dispersers, protects cells from photooxidative damage by absorbing high-energy light, and alleviates oxidative stress by scavenging free radicals [15]. However, in turfgrasses, the purple color of stolons and seed heads can affect the aesthetics of an otherwise homogeneous green lawn. The genetic loci responsible for anthocyanin production in various organs in zoysiagrass cultivars have yet to be identified. A comparative study on purple *versus* green spikes of two *Z. japonica* cultivars showed that the genes encoding dihydroflavonol 4-reductase (*DFR1)* and anthocyanidin synthase (*ANS1)* were significantly upregulated in purple-spiked plantsAhn, Kim [8]. However, both genes lacked variation in the coding region as well as ∼450 bp of upstream sequence between the anthocyanin-producing and green accessions, suggesting that the differential regulation of these genes may be governed by a transcription factor [8].

The primary aim of this study was to develop a high-density linkage map in a biparental F_2_ population derived from a cross between a commercial *Z. japonica* cultivar, Meyer, and a *Z. matrella* Plant Introduction (PI), PI 231146, as a tool to determine the genetic mechanisms underlying traits, particularly those that differ between the two species. Although the two parents and the F_1_ progeny selfed to produce the F_2_ population had purple stolons and seed heads, the F_2_ progeny segregated for anthocyanin production in both organs. Seed head color was segregating in an F_1_ population generated from the same parents. Therefore, the secondary aim of our study was to map the genomic regions associated with seed head and stolon pigmentation in both the newly generated F_2_ linkage map and in existing parental F_1_ maps [16], and identify the causal genes. Variation in anthocyanin pigmentation mapped to the same locus, *PP*, in both the F_2_ and F_1_ maps. We identified two mutations in a MYC-bHLH transcription factor that regulates anthocyanin biosynthesis, with one mutation being heterozygous in Meyer, and the second being heterozygous in PI 231146. The SNP variant present in Meyer affects both stolon and seed head color while the SNP variant in PI 231146 affects only seed head color. Understanding the regulatory genes controlling anthocyanin biosynthesis is essential for zoysiagrass breeders aiming to enhance turfgrass aesthetic value. Further, interspecific genetic maps are invaluable tools for mapping traits that differentiate species and will play a crucial role in advancing breeding efforts.

## Results

### Genetic map

The number of paired-end reads obtained through genotyping-by-sequencing (GBS) averaged 2 638 279 per sample with a range of 1 572 677 to 5 028 129. A total of 48 443 SNPs were called using Haplotype Caller within the Genome Analysis Toolkit (GATK) [17], and 50 133 SNPs using GATK’s Unified Genotyper. Filtering for markers that were heterozygous in F_1_-19-TZ-14321, the F_1_ that was selfed to generate the F_2_ population, exhibited less than 20% missing data across the F_2_ progeny (with per sample read depth ≥ 8x), and with chi-squared p-values > 1 x 10^-10^ for 1:2:1 segregation resulted in 7 698 SNPs from Haplotype Caller and 8 725 SNPs from Unified Genotyper. Merging of the two SNP sets yielded 10 283 non-redundant SNPs of which 1 559 (15.2%) were Haplotype Caller-specific, 2 587 (25.2%) were Unified Genotyper-specific, and 6 137 (59.7%) were common SNPs.

Duplication followed by reversing in one copy the scores of markers for which the parental genotypes were ambiguous (C or D) or unknown increased the number of SNPs to 14 128. Reducing sets of cosegregating markers to a single representative marker decreased the dataset to 10 895 SNPs, and these were input into MST-map. At LOD 12, 32 linkage groups were obtained. Removal of linkage groups consisting only of duplicate markers left 22 linkage groups for input into SeSAM’s autoMap function for marker ordering. A total of 89 markers exhibiting more than 10 double recombination events across the 530 progenies were removed from the maps. Due to severe segregation distortion (chi-squared p-values < 1x10^-10^ for 1:2:1 segregation) in the distal regions of chromosomes 2 and 7, and central regions of chromosomes 5 and 19, markers derived from these regions had been removed at the start of the mapping process, which led to incomplete linkage groups. After incorporating 753 highly distorted markers into these four chromosomes, the final map comprised 8 402 SNPs organized into 20 linkage groups (Fig. 1; Table S1). On average, each linkage group spanned 89 cM, with lengths ranging from 53.5 to 126.4 cM, and contained an average of 420 markers (range: 124–832 markers) (Table 1). The average distance between markers was 0.21 cM with only two intervals across the 20 chromosomes ≥ 5 cM. Segregation distortion was observed in multiple chromosomes, mostly caused by an overrepresentation of B alleles (Table S1; Fig. S1). Linkage groups (LGs) were named in accordance with the *Z. japonica* pseudomolecules with each even number and the subsequent uneven number (*e.g.* chromosomes 1 and 2, 3 and 4) representing homoeologous chromosomes [4].

**Figure 1.**
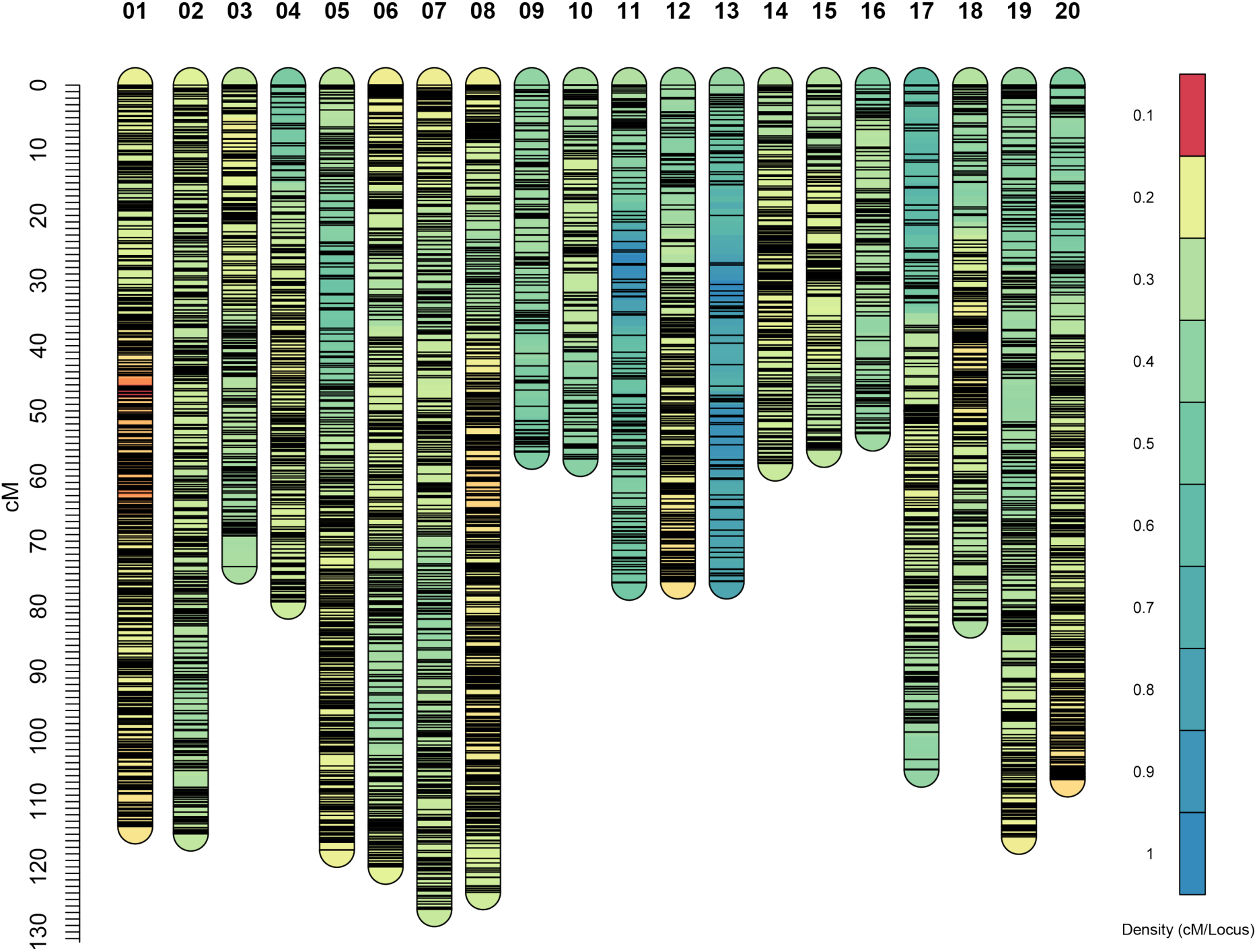
High density linkage map generated in a *Z. japonica* acc. Meyer x *Z. matrella* acc. PI 231146 F_2_ population. The map comprises 8 402 SNP markers across the 20 zoysiagrass chromosomes. Marker names and locations are provided in Table S1.

**Table 1.** Map statistics.

| Zoysiagrass F<sub>2</sub> linkage maps |
| --- |

| Chromosome | Chromosome length in genome assembly (bp) | # Markers | Length (cM) | Position of first marker on genome assembly | Position of last marker on genome assembly | Region covered by LG (Mb) | % coverage |
| --- | --- | --- | --- | --- | --- | --- | --- |
| 1 | 18 983 185 | 832 | 113.7 | 17 062 | 18 967 069 | 18 950 007 | 99.8% |
| 2 | 17 914 192 | 537 | 114.8 | 7 724 | 17 912 629 | 17 904 905 | 99.9% |
| 3 | 8 228 935 | 352 | 73.9 | 53 699 | 8 188 299 | 8 134 600 | 98.9% |
| 4 | 11 297 323 | 393 | 79.3 | 91 675 | 11 270 996 | 11 179 321 | 99.0% |
| 5 | 19 970 953 | 556 | 117.3 | 24 704 | 19 402 653 | 19 377 949 | 97.0% |
| 6 | 19 679 654 | 593 | 119.9 | 11 604 | 19 662 421 | 19 650 817 | 99.9% |
| 7 | 20 256 391 | 591 | 126.4 | 30 989 | 20 254 110 | 20 223 121 | 99.8% |
| 8 | 21 132 631 | 800 | 123.8 | 42 234 | 20 550 225 | 20 507 991 | 97.0% |
| 9 | 8 250 627 | 169 | 56.3 | 28 785 | 8 236 420 | 8 207 635 | 99.5% |
| 10 | 8 173 407 | 235 | 57.4 | 11 961 | 8 106 371 | 8 094 410 | 99.0% |
| 11 | 11 083 367 | 175 | 76.4 | 40 210 | 11 082 329 | 11 042 119 | 99.6% |
| 12 | 14 849 823 | 415 | 76.2 | 59 344 | 14 818 153 | 14 758 809 | 99.4% |
| 13 | 11 692 151 | 124 | 76.2 | 17 912 | 11 596 382 | 11 578 470 | 99.0% |
| 14 | 8 885 846 | 320 | 58.1 | 23 073 | 8 431 857 | 8 408 784 | 94.6% |
| 15 | 7 690 864 | 289 | 56.0 | 45 077 | 7 655 559 | 7 610 482 | 99.0% |
| 16 | 8 056 988 | 199 | 53.5 | 44 312 | 8 044 728 | 8 000 416 | 99.3% |
| 17 | 10 678 808 | 390 | 105.1 | 9 100 | 10 664 610 | 10 655 510 | 99.8% |
| 18 | 11 179 804 | 423 | 82.2 | 22 588 | 11 176 547 | 11 153 959 | 99.8% |
| 19 | 19 485 313 | 471 | 115.4 | 78 773 | 19 268 726 | 19 189 953 | 98.5% |
| 20 | 16 245 370 | 538 | 106.5 | 93 681 | 16 240 059 | 16 146 378 | 99.4% |
| Total | 273 735 632 | 8 402 | 1 788.4 |  |  | 270 775 636 |  |
| Average |  | 420.10 | 89.0 |  |  |  | 98.9% |

Alignment of the GBS reads to the *Z. japonica* scaffold assembly (ZJN_r.1.1.fa) combined with BLASTN results using 1-kb of *Z. japonica* scaffold sequence surrounding the mapped markers as queries against the *Z. japonica* pseudomolecules identified the scaffolds from the ZJN_r.1.1.fa assembly corresponding to each pseudomolecule (Tables S1, S2). This analysis revealed that the linkage maps included markers derived from scaffolds that were not integrated into the pseudomolecule assembly (ZJN_pseudomol.r1.0.fa) and also identified chimeric scaffolds (Table S2). The linkage groups covered, on average, 98.9% of the chromosomes (range 94.6% – 99.9%) (Table 1).

BLASTN analyses with the same 1-kb of marker-associated *Z. japonica* sequence against the *Z. matrella* and *Z. pacifica* scaffold-level genome assemblies consistently identified two highly similar top hits in *Z. matrella*, compared to one in *Z. pacifica* (Table S1). This is likely due to the high level of heterozygosity present in *Z. matrella*, potentially due to a *Z. japonica* x *Z. pacifica* hybrid origin [3, 18] causing scaffolds for both haplotypes to be represented in the assembly. The complete linkage maps with genotypic scores and comparative relationships to the *Z. japonica*, *Z. matrella* and *Z. pacifica* genome assemblies are provided in Table S1.

### Comparative analyses of the zoysiagrass F_2_ linkage map with the finger millet genome

Comparative analysis of the zoysiagrass F_2_ linkage map with the KNE 796-S v1.1 genome assembly [19] of its chloridoid relative, finger millet, showed a highly syntenic relationship with each pair of homoeologous *Zoysia* chromosomes largely corresponding to a pair of homoeologous finger millet chromosomes (Fig. 2A; Table S2). For simplicity, the *Zoysia* relationship to only finger millet subgenome A is shown in Fig. 2A. Colinearity was highly conserved for most *Zoysia* – finger millet syntenic pairs (Fig. S2; Table S1). The exceptions were the homoeologous zoysiagrass chromosomes 9 (Chr09) and 10 (Chr10), which had both a lower percentage of comparative markers than other chromosomes and an overall lower level of colinearity (Fig. S2; Table S3). Several *bona fide* inversions of a few megabases that occurred in zoysiagrass were observed relative to finger millet in otherwise highly colinear chromosomes, including on zoysiagrass Chr08, Chr17 and Chr20. These rearrangements were typically present in only one of the two homoeologous *Zoysia* chromosomes (Fig. S2). The comparative relationships also showed that zoysiagrass chromosomes 11 and 12, like its finger millet orthologs 4A and 4B, are acrocentric (Fig. S2), a configuration that likely originated before the radiation of the grasses [20].

**Figure 2.**
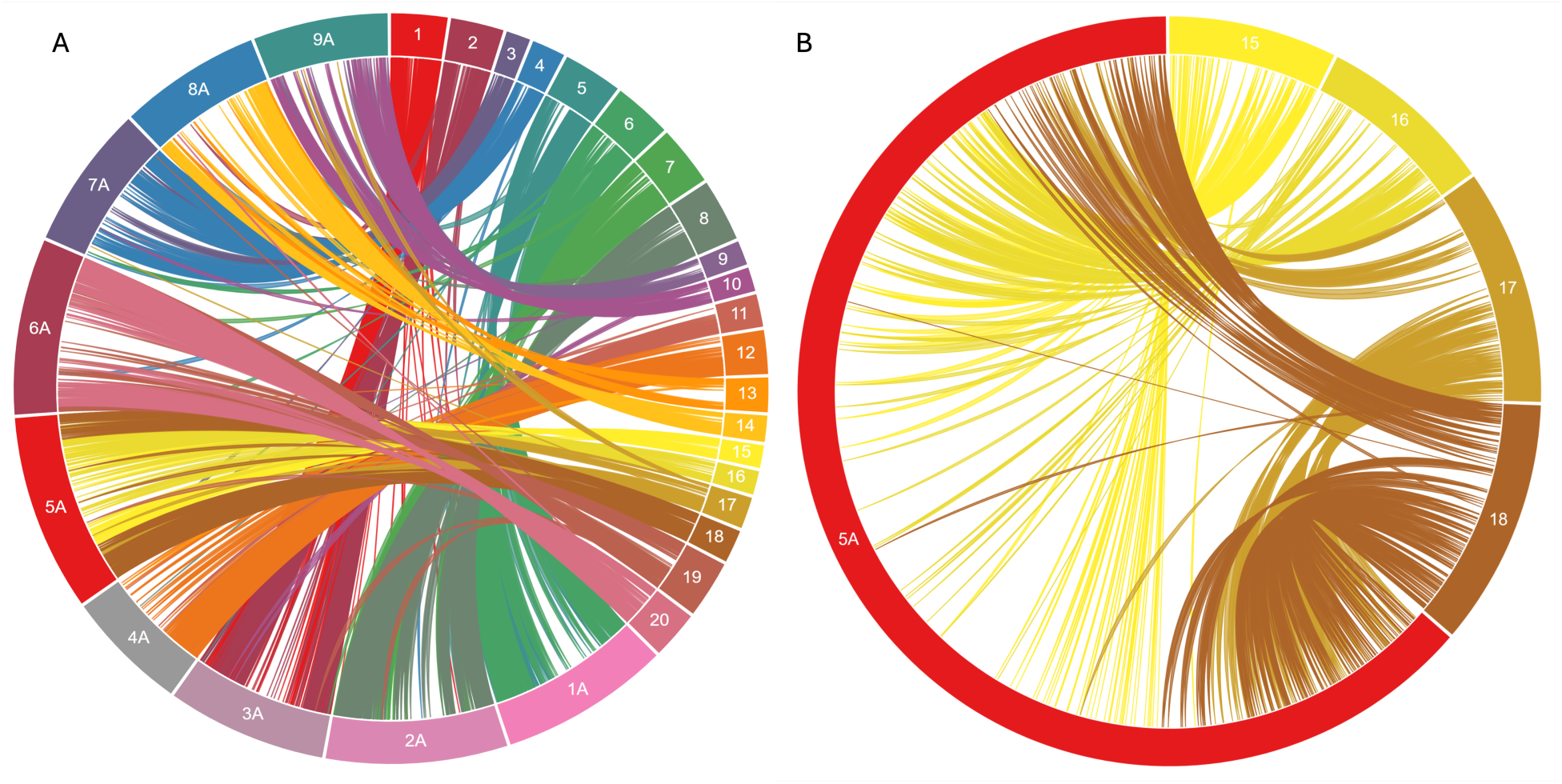
Circos diagrams showing comparative relationships. (A) Relationship between the Meyer x PI 231146 F_2_ linkage map and the *Eleusine coracana* (finger millet) KNE 796-S genome assembly. For simplicity, only the finger millet A genome is shown. (B) Syntenic relationships between *Eleusine coracana* chromosome 5A and *Zoysia japonica* pseudomolecules 15, 16, 17, and 18. A chromosome reduction event from *n* = 10 to *n* = 9 that gave rise to the genus *Eleusine* resulted in chromosome 5 exhibiting synteny with two distinct pseudomolecules in *Zoysia*. Figures were generated using CIRCA (OMGenomics, http://omgenomics.com/circa/).

Both zoysiagrass (2*n* = 4*x* = 40) and finger millet (2*n* = 4*x* = 36) are allotetraploids, but the lineage leading to finger millet underwent a chromosome reduction through insertional dysploidy [19] from *x* = 10, the ancestral basic chromosome number in the Chloridoideae, to *x* = 9. Therefore, one pair of finger millet chromosomes, 5A and 5B, is syntenic to two pairs of zoysiagrass chromosomes, 15/16 and 17/18 (Fig. 2B, Fig. S2). *Zoysia* chromosomes 17 and 18 are orthologous to the distal regions of finger millet chromosomes 5A and 5B, and *Zoysia* chromosomes 15 and 16 to the proximal regions of finger millet 5A and 5B.

### QTL for seed head and stolon pigmentation

The two parents used in this study, Meyer and PI 231146, both exhibit purple pigmentation in their stolons and seed heads. While F_1_-19-TZ-14321, the F_1_ that was selfed to generate the F_2_ population, also has purple pigmentation in stolons and seed heads, the F_2_ population is segregating 1:3 (green:purple) for both traits, indicating that the trait is controlled by a single gene (Fig. 3A-D; Table 2; Table S4). The F_1_ plant and either one or both parents must therefore be heterozygous for anthocyanin production. Stolon and seed head color largely cosegregate in the F_2_ progeny and mapped to a single region on chromosome 12 (Fig. 3E). The large-effect QTL explained between 68% and 89% of the phenotypic variation (Table 3). Despite the cosegregation of both traits in the F_2_ population, F_1_ progeny derived from the same cross uniformly had purple stolons (three lines with green stolons were likely off-types and were removed from the analysis) but segregated 1:3 (green: purple) for seed head color (Table 2; Table S5), indicating that the two traits are controlled independently. QTL analyses conducted on the maternal (Meyer) and paternal (PI 231146) F_1_ genetic maps [16] identified overlapping QTL on chromosome 12 (Fig. S3). The QTL regions identified in the F_1_ map also overlapped with the QTL identified in the F_2_ population (Table 3).

**Figure 3.**
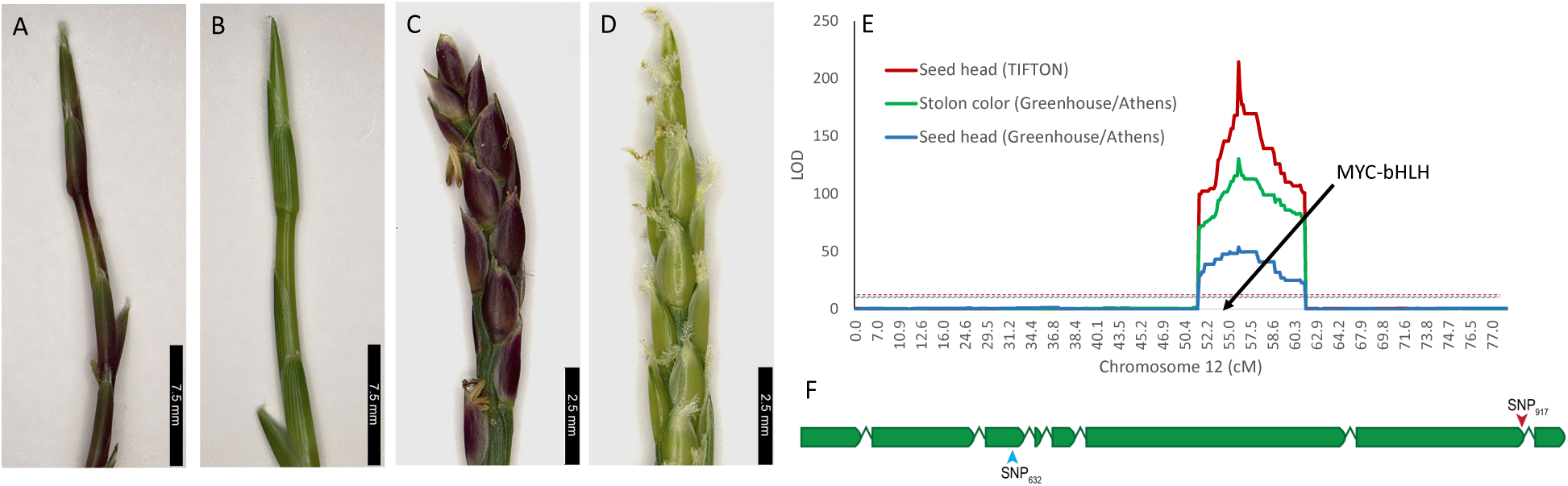
Anthocyanin presence in stolons and seed heads, QTL locations for both traits and schematic of causal gene. Representative images showing anthocyanin presence (A, C) in stolons (A, B) and seed heads (C, D) in F_2_ progeny from the Meyer x PI 231146 cross. (E) Quantitative trait loci for anthocyanin pigmentation on chromosome 12. LOD: logarithm of the odds. Black arrow points to the position of the *PP* gene (MYC-bHLH). (F) Schematic representation of the *PP* gene (*Zjn_sc00004.1.g07010.1.sm.mk)* on chromosome 12. Introns are not drawn to scale. The SNP at position Zjn_sc00004.1:2 965 917 (SNP_917_) that leads to intron retention is indicated with a red arrowhead. The SNP at position Zjn_sc00004.1:2 967 632 (SNP_632_) is indicated with a blue arrowhead.

**Table 2.** Phenotypic data summary of anthocyanin presence (purple) and absence (green) in F_1_ and F_2_ population in different tissue types.

| Population and Tissue type<sup>1</sup> | Green | Purple | Ratio | p-value |
| --- | --- | --- | --- | --- |
| F<sub>2</sub> stolon color (Greenhouse/ Athens) | 136 | 371 | 1:3 | 0.343 |
| F<sub>2</sub> seed head color (Greenhouse/ Athens) | 34 | 103 | 1:3 | 0.961 |
| F<sub>2</sub> seed head color (Nursery/Tifton) | 101 | 324 | 1:3 | 0.556 |
| F<sub>1</sub> stolon color (Tifton) | 3<sup>2</sup> | 155 | - |  |
| F<sub>1</sub> seed head color (Tifton) | 44 | 114 | 1:3 | 0.408 |

<sup>1</sup> Replicated plants from the F<sub>2</sub> population were grown in a greenhouse in Athens, GA, and a nursery at Tifton, GA. The F<sub>1</sub> population was grown in a nursery in Tifton, GA
<sup>2</sup> Likely off-types (not further analyzed)

**Table 3.** Quantitative trait loci (QTL) for anthocyanin variation in seed heads and stolons.

| | Number<br>of<br>progeny<br>pheno-<br>typed | Chro<br>moso<br>me | Marker at<br>QTL peak | Position<br>(cM) of<br>QTL<br>peak | Position<br>(bp) of QTL<br>on pseudo-<br>molecule | Marker at<br>left border<br>of QTL<br>region | Position<br>(cM) of<br>left<br>border<br>marker | Position<br>(Mb) of<br>left border<br>marker on<br>pseudo-<br>molecule | Marker<br>at right<br>border of<br>QTL<br>region | Position<br>(cM) of<br>right<br>border<br>marker | Position<br>(Mb) of<br>right border<br>marker on<br>pseudo-<br>molecule | LOD<br>at<br>QTL<br>peak | LOD<br>threshold<br>($\alpha < 0.05$) | PVE<br>% |
| --- | --- | --- | --- | --- | --- | --- | --- | --- | --- | --- | --- | --- | --- | --- |
| **F<sub>2</sub> population<sup>1</sup>** |  |  |  |  |  |  |  |  |  |  |  |  |  |  |
| Seed head color<br>(Tifton Nursery) | 425 | 12 | Tag_5328 | 55.2 | 11 600 003 | Tag_5211r | 50.2 | 11 102 906 | Tag_5513 | 59.7 | 12 482 465 | 213.7 | 4.3 | 89 |
| Stolon color<br>(Athens GH) | 507 | 12 | Tag_5328 | 55.2 | 11 600 003 | Tag_5211r | 50.2 | 11 102 906 | Tag_5513 | 59.7 | 12 482 465 | 127.9 | 4.3 | 68 |
| Seed head color<br>(Athens GH) | 137 | 12 | Tag_5328 | 55.2 | 11 600 003 | Tag_5211r | 50.2 | 11 102 906 | Tag_5513 | 59.7 | 12 482 465 | 54.1 | 5.6 | 81 |
| **F<sub>1</sub> population<sup>1</sup>** |  |  |  |  |  |  |  |  |  |  |  |  |  |  |
| Seed head<br>(Tifton) in<br>paternal map | 148 | 12 | Tag_683r | 63.0 | 11 457 570 | Tag_666 | 57.5 | 10 887 884 | Tag_712 | 66 | 12 223 422 | 19.3 | 4.1 | 45 |
| Seed head<br>(Tifton) in<br>maternal map | 148 | 12 | Tag_675r | 38.4 | 11 164 131 | Tag_671 | 37.5 | 11 102 906 | Tag_685 | 45 | 11 557 157 | 6.8 | 4.1 | 15 |

<sup>1</sup> Anthocyanin variation was scored in F<sub>1</sub> and F<sub>2</sub> populations derived from a cross between Meyer and PI 231146, maintained either in a greenhouse
(GH) in Athens, GA or in a nursery in Tifton, GA

### Purple coloration is controlled by a MYC-bHLH transcription factor

The QTL identified in the F_2_ population was projected on the finger millet KNE 796-S A genome (Chr4A, ∼7 400 000 bp to ∼7 900 000 bp), and the 319 annotated genes located in the finger millet region orthologous to the zoysiagrass QTL were extracted (Table S6). Two genes, *ELECO.r07.4AG0307750* and *ELECO.r07.4AG0307030*, had gene descriptions associated with the term ‘anthocyanin’ (Table S6). No ortholog of *ELECO.r07.4AG0307030* was found in the QTL region on chromosome 12 in zoysiagrass, although an ortholog was identified in the expected region on the homoeologous zoysia chromosome 11. Further, *ELECO.r07.4AG0307030* had the highest homology in NCBI’s SwissProt database to homeobox-leucine zipper protein RICE OUTERMOST CELL-SPECIFIC 4 (ROC4), a transcription factor controlling the production of cuticular wax and bulliform cell shrinkage [21]. The most likely causal gene to the QTL therefore was the zoysia ortholog of *ELECO.r07.4AG0307750*, which encodes a MYC-bHLH transcription factor that regulates anthocyanin production in anthers and stigma in finger millet [19] and is orthologous to the maize anthocyanin regulatory gene *R*-S (*Seed color component at R1*; Genbank acc. P13027.1) [22]. The *Z. japonica* ortholog, *Zjn_sc00004.1.g07010.1.sm.mk*, was located ∼30 kb from the QTL peak (Fig. 3E), and will be referred to as *PP* following the study in finger millet [19]. Although *Z. japonica* and *Z. matrella* are tetraploids, *Zjn_sc00004.1.g07010.1.sm.mk* does not have a homoeolog in either species, so its inactivation is sufficient to eliminate anthocyanin production.

Whole genome Illumina sequencing of Meyer and PI 231146, the parents of the mapping population, identified five missense mutations that were homozygous for the reference allele in Meyer and heterozygous in PI 231146, one SNP that was heterozygous in both Meyer and PI 231146, and one SNP that was homozygous for the reference allele in PI 231146 and heterozygous in Meyer (Table S7). The latter SNP (position 2 965 917 bp on Zjn_sc00004.1), which we will refer to as SNP_917_, was located at the very end of the seventh exon in Meyer (Fig. 3F). Analysis of RNA-seq reads (SRA DRA001679) obtained from *Z. japonica* lines with green and purple seed heads [8] indicated that the G → A mutation at SNP_917_ was present in the green line and resulted in intron retention, likely rendering the protein non-functional (Fig. S4). Intron retention in *Zjn_sc00004.1.g07010.1.sm.mk* was experimentally validated by amplifying across intron 7 in Meyer, PI 231146, two purple F_2_ progeny and two green F_2_ progeny, and sequencing of the two fragment sizes (Fig. 4). Genotyping of a subset of the F_2_ population with a *Dde*I cleaved amplified polymorphic sequence (CAPS) marker specific for the G → A substitution at SNP_917_ in exon 7 showed that SNP_917_ was either homozygous GG or heterozygous GA in all purple progeny tested (n = 16), and homozygous AA in all green progeny tested (n = 13) (Table S8).

**Figure 4.**
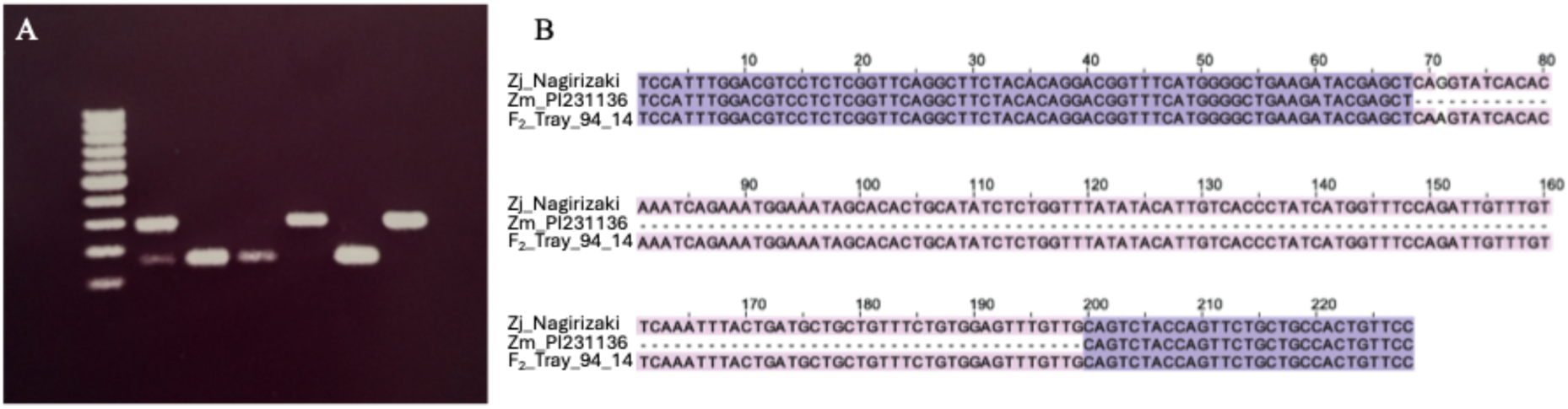
Retention of intron 7 in *Zjn_sc00004.1.g07010.1.sm.mk*. (A) Gel image showing amplification across the intron 7 region of *Zjn_sc00004.1.g07010.1.sm.mk*. Lanes (from left): 100 bp DNA ladder, Meyer (heterozygous G/A for the G → A SNP at the end of exon 7 (SNP_917_); purple), PI 231146 (homozygous GG; purple), F_2_ progenies Tray_14_12 (homozygous GG; purple), Tray_29_4 (homozygous AA; green), Tray_19_12 (homozygous GG; purple) and Tray_87_13 (homozygous AA; green). The top fragments represent amplicons with intron 7 retained, and bottom fragments represent amplicons with intron 7 excised. Presence of both bands signifies a heterozygous SNP. (B) Sequence of amplicons derived from progeny Tray_94_14 (genotype AA; green stolons and seed head) and parental line PI 231146 (genotype GG; purple stolons and seed heads), and corresponding sequence from *Z. japonica* Nagirizaki genome assembly.

Because all F_1_ progeny from the Meyer x PI 231146 cross had purple stolons but segregated for seed head color, we assayed 153 F_1_ plants with the *Dde*I CAPS marker. All F_1_ tested were either homozygous GG or heterozygous GA, consistent with purple stolon coloration (Table S5). Co-mapping of the QTL for seed head color in the F_1_ population with the QTL for seed head/stolon color in the F_2_ population suggested that there was a second mutation either in the same gene (hypothesis 1) or in a nearby gene (hypothesis 2). Under hypothesis 1 (Fig. S5A), the second mutation in *MYC-bHLH* would be expected to be heterozygous in PI 231146 considering that the intron-retention SNP in *MYC-bHLH* was heterozygous in Meyer and that both parents had purple seed heads. Further, F_1_ progeny heterozygous for both mutations would be expected to have green seed heads. If, however, the mutation controlling seed head color was in a different gene in the same region (hypothesis 2), green seed heads would require plants to be homozygous for the second mutation (Fig. S5B). To test hypothesis 1, we genotyped a subset of the F_1_ plants for the G → T SNP at Zjn_sc00004.1:2 967 632, which we will refer to as SNP_632_ (Table S5). SNP_632_ is one of three missense variants that are heterozygous in PI 231146, homozygous wild type in Meyer, and homozygous wild type in F_1_-19-TZ-14321, the F_1_ that gave rise to the F_2_ population (Table S7). Homozygosity for the wild type allele in F_1_-19-TZ-14321 would be expected under scenario 1 because the F_1_ is heterozygous for the exon 7 mutation and has purple seed heads. SNP_632_ leads to an Alanine (Ala) to Serine (Ser) amino acid substitution in the N-terminal region of MYC-bHLH (Fig. S6). Assaying SNP_632_ with a *Bss*HII CAPS marker revealed that all F_1_ plants tested with green seed heads (n = 41) were heterozygous GT, while F_1_ with purple seed heads (n = 108) were either heterozygous GT or homozygous GG (Table S5). However, plants heterozygous for GT displayed seed head anthocyanins only if homozygous GG (wild type) at location SNP_917_ (exon 7 SNP). On the other hand, all plants with green seed heads were heterozygous at both loci.

### Yeast two-hybrid assays demonstrate functional effect of SNP_632_

To test whether the Ala to Ser substitution caused by SNP_632_ affects the binding of MYC-bHLH with MYB transcription factors, we conducted yeast two-hybrid assays using the N-terminal region of MYC-bHLH_Ala_ and MYC-bHLH_Ser_ as bait and three *Z. japonica* MYB transcription factors most closely related to maize C1 (encoded by *Z. japonica* acc. Nagirizaki genes with IDs *Zjn_sc00004.1.g07010.1.sm.mk, Zjn_sc00049.1.g03240.1.am.mk* and *Zjn_sc00010.1.g01550.1.am.mk*; Fig. S7) as prey. The assays indicated that the strength of binding varied by MYB factor. Further, the Ala to Ser substitution present in MYC-bHLH affected the interaction with MYB, but this effect was not equal for all MYB factors (Fig. 5). The amino acid substitutions present in MYBb and MYBc had no discernable effect on binding.

**Figure 5.**
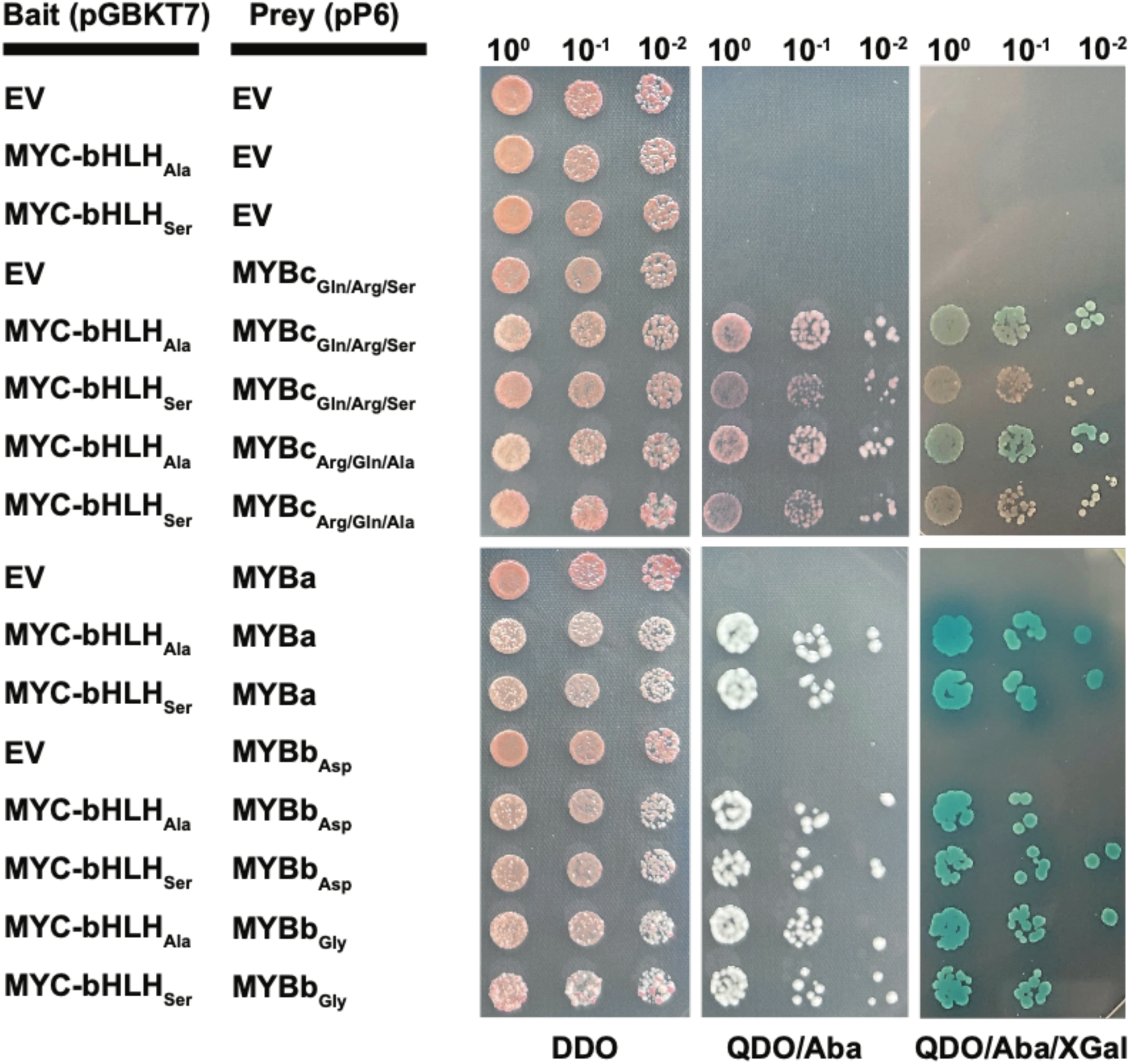
Physical interactions between MYC-bHLH and MYB transcription factors analyzed using yeast two-hybrid assays. Representative colonies of *S. cerevisiae* expressing bait and prey fusion protein pairs. MYC-bHLH_Ala_ and MYC-bHLH_Ser_ were cloned into the bait vector pGBKT7. MYBa, MYBb and MYBc were cloned into the prey vector pP6. Two MYBb variants that differed by a single amino acid, MYBb_Asp_ and MYBb_Gly_, and two MYBc variants that differed by three amino acids, MYBc_Gln/Arg/Ser_ and MYBc_Arg/Gln/Ala_ were assayed. Successful co-transformation of prey and bait was assessed on DDO medium (SD/–Leu/–Trp) and interactions were assessed on QDA/Aba (SD/–Leu/–Trp /–His/–Ade + Aureobasidin A), and QDO/Aba/ XGal (SD/–Leu/–Trp /–His/–Ade + Aureobasidin A + X-α-Gal) media. EV indicates empty vector controls.

## Discussion

### Zoysiagrass high-density genetic map

The SNPs in this study were called using GATK [17] with both Unified Genotyper and Haplotype Caller. Use of Haplotype Caller is recommended and Unified Genotyper only exists in version 3.2 of GATK for legacy reasons. However, a previous study had found that, in general, Unified Genotyper identified more mappable, and hence good quality, SNPs than Haplotype Caller [23]. To maximize the number of markers for genetic mapping, both SNP calling functions were used in the current study, and SNPs were merged by position to create a non-redundant set of markers for genetic map construction. Ultimately, 25.3% of the SNPs included in the high-density map were identified only by Unified Genotyper, 12.5% only by Haplotype caller, and 62.2% by both. These percentages are comparable to those obtained by Qi and colleagues [23] in finger millet in which 20% of mapped SNPs were called only by Unified Genotyper and 12% only by Haplotype Caller, and reinforce the value of using Unified Genotyper solely or combined with Haplotype Caller to maximize SNP identification.

We used a hybrid approach, capitalizing on the strength and speed of different mapping programs, to generate robust linkage maps. MSTMap, based on the traveling salesman principle, is highly efficient to identify linkage groups but does not always give robust marker orders. SeSAM uses a seriation and placement-based approach to make a robust framework map, and then places the remaining markers in relation to the framework map. In a last step, we used a modified version of MAPMAKER [23, 24] to remove markers with more double recombination events than a chosen threshold (∼2% in this study), and to calculate the final map distances in Kosambi centiMorgans. Marker orders were manually verified against the *Z. japonica* genome assembly, and markers not separated by clear recombination events were reordered in accordance with marker orders on the genome assembly. No reordering was done if this led to an increase in recombination events. The resulting robust high-density genetic map represents the first interspecific *Z. japonica* x *Z. matrella* F_2_ map generated for zoysiagrass. Genetic maps for zoysiagrass have been previously reported [5, 6, 25], including one from the same set of parents [16], but have been limited to mapping intraspecies recombination. An interspecific F_2_ population design is key to mapping traits that differentiate the two species.

A total of 8 402 SNP markers were organized into 20 linkage groups. Tanaka et al. [4] provided both scaffold-level and pseudomolecule-level assemblies for *Z. japonica*. Linkage maps are important tools for enhancing genome assemblies by resolving errors and filling gaps. Accordingly, we identified scaffolds in multiple chromosome regions that were not incorporated in the pseudomolecule-level assembly (Table S2). We also identified 16 chimeric scaffolds (seven in addition to the nine that were reported by Tanaka et al. [4]) (Table S2), and at least 16 scaffolds that were incorporated in an incorrect position or in inverse orientation in the pseudomolecules (Table S1). Scaffolds included in the *Z. japonica* pseudomolecule assembly but absent from the genetic map are also listed in Table S2.

The genetic map reflects recombination between *Z. japonica* and *Z. matrella* chromosomes. *Z. matrella* is thought to have a hybrid *Z. japonica* x *Z. pacifica* origin [3, 18], and the higher diversity between the haplotypes likely impeded their collapse in the *Z. matrella* genome assembly [4]. The presence of twice the expected gene copies led Wang et al. [18] to conclude, incorrectly, that *Z. matrella* had undergone a species-specific whole-genome duplication event following its divergence from *Z. pacifica*. As our maps demonstrate, *Z. japonica* and *Z. matrella* are fully cross-compatible. While there is a two-fold difference in genetic length across the 20 *Zoysia* chromosomes (Fig. 1), the significant positive correlation between the genetic map length and physical chromosome length (r^2^ = 0.85; p <0.0001; Fig. S8) suggests that the variation in recombination is not caused by variation in the ratio of *Z. japonica* and *Z. pacifica* alleles across *Z. matrella* chromosomes, assuming a hybrid origin for *Z. matrella*.

We do, however, observe very severe segregation distortion in several of the linkage groups. In most cases, distortion strongly favored the Meyer (female) allele, particularly in the distal regions of chromosomes 2 and 7, and the centromeric regions of chromosomes 5 and 19 (Fig. S1). Segregation distortion can have many underlying causes, including the presence of a killer (derived from Meyer) – target (derived from PI 231146) drive system in which gametes carrying the target element are destroyed during or following meiosis, or a poison – antidote (both derived from Meyer) drive system in which gametes lacking the antidote are rendered unviable [26]. The killer and target loci, and the poison and antidote loci, should be closely linked to avoid self-destruction, and would expected to be located in the region displaying the highest segregation distortion. Transmission bias was complete for the loci on chromosome 5 and chromosome 7 because only Meyer alleles (∼33%) and heterozygotes (∼66%) were identified. Bias against the Meyer allele was also observed in a few regions, most notably on chromosome 18. This highlights the complexity of allele dynamics in this interspecific population, which has ramifications for breeding. Further work is needed to determine whether the observations made in the Meyer x PI 231146 cross apply more widely to interspecific *Z. japonica* x *Z. matrella* crosses, and whether the direction of the cross has an effect [27].

Although homoeologous chromosomes have been identified in zoysiagrass linkage maps using comparative information ([5]; this study) and in the *Z. japonica* pseudomolecules, the two subgenomes of zoysiagrass remain unidentified. Jia and colleagues [28] were unable to phase the two subgenomes in *Z. japonica* using SubPhaser. We tested polyCRACKER [29], which performs unsupervised partitioning of polyploid subgenomes by analyzing signatures of repetitive DNA evolution, but were similarly unsuccessful (File S1). The divergence age between the zoysiagrass subgenomes has been estimated at ∼20.8 million years ago (MYA)[18]. However, the age of the allopolyploidization event is unknown. Considering the inability of software such as SubPhaser [28] and polyCRACKER [29], which rely on subgenome-specific repeats, to partition the chromosomes into two subgenomes, it seems likely that the hybridization leading to tetraploid zoysiagrass is not a recent event. Transposable elements (TEs) are removed over time through homologous or illegitimate recombination [30, 31]. Therefore, the majority of TEs present in a genome will likely have inserted in the last eight million years or so [32]. If allopolyploidization occurred more than 10 million years ago, the majority of subgenome-specific repeats will have eroded. Further, transposable elements that have been active post-polyploidization will typically not display subgenome preference, abolishing their use in subgenome identification. Identification of the subgenome chromosomes will likely require information on at least one of the diploid progenitors or on a species that is taxonomically more closely related to one of the diploid progenitors [33].

### Synteny between zoysiagrass and finger millet

Colinearity is highly conserved between the genetic map of *Zoysia* and the genome assembly of its chloridoid relative, finger millet (Fig. 2; Fig. S2). The exceptions are *Zoysia* Chr09 and Chr10 which are syntenic to finger millet chromosomes 9A and 9B. The orthologous rice (Chr11), foxtail millet (Chr08), switchgrass (Chr08) and sorghum (Chr05) chromosomes all have the highest abundance of disease resistance genes [34–38], and it is likely that this is also the case for zoysiagrass Chr09 and Chr10, and finger millet chromosomes 9A and 9B. This high concentration of disease resistance genes likely drives rapid adaptive divergence through an arms race with pathogens, leading to frequent shifts in genomic structure and an overall lower level of colinearity.

### Two alleles of the MYC-bHLH transcription factor affect anthocyanin loss in different organs

Anthocyanins are the major plant flavonoid compounds, which confer appealing colors to flowers, fruits and other organs, and contribute to stress tolerance. In zoysiagrass, anthocyanins are found in both stolons and seed heads (Fig. 3), specifically the glumes, stigma and anthers. The anthocyanin biosynthesis genes are activated as a single unit by a ternary complex of the MYB-bHLH-WD40 transcription factors (MBW complex) [39–41]. Functional inactivation of the MYC-bHLH transcription factor in the MBW complex has previously been shown to be causal to the loss of anthocyanins in anthers and stigma of finger millet [19]. The single large-effect QTL identified in our study for the variation in stolon and seed head coloration encompassed the finger millet *PP* region that carries the MYC-bHLH anthocyanin regulatory gene *ELECO.r07.4AG0307750*, making the zoysiagrass ortholog, *Zjn_sc00004.1.g07010.1.sm.mk*, a strong candidate gene for the QTL. Green coloration at the QTL locus was associated with homozygosity of the A (Meyer) allele, indicating that Meyer, which has purple stolons and seed heads, is heterozygous for the mutation. Two variants located in *Zjn_sc00004.1.g07010.1.sm.mk* were heterozygous in Meyer as well as in the F_1_ that was selfed to generate the F_2_ population, and homozygous (SNP_917_) or heterozygous (SNP_730_) in PI 231146 (Table S7). SNP_917_, which is the last base of exon 7, leads to retention of intron 7 (Fig. 4; Fig. S4). RNA splicing depends on conserved intron-exon sequences. In addition to the almost completely conserved “GT” splice donor site and “AG” splice acceptor site, consensus sequences have also been identified immediately up- and downstream of introns in the neighboring exons. In grasses, the last base of an exon is G in more than 75% of the cases while A occurs at a frequency of ∼10% [42]. The G → A mutation at position 2 965 917 alters the exon boundary and impairs splice site recognition, leading to intron retention and a protein with an altered C-terminus that lacks part of the MYC-bHLH ACT-like domain (Fig. S6). The ACT-like domain, as demonstrated in the maize bHLH transcription factor R, is critical for regulating anthocyanin biosynthesis [43]. In the presence of the ACT domain, the bHLH domain remains in monomer form and interacts with the R-interacting factor 1 (RIF1) to bind to the promoter of a subset of the anthocyanin biosynthesis genes through interaction with the MYB factor C1 [44]. In the absence of ACT, the bHLH domain dimerizes and gains capability to bind to G-box motifs but interaction with RIF1, which is essential for anthocyanin production by R and C1 in maize, is lost [44, 45]. In addition, the alteration of the C-terminus of the ACT domain by the retention of intron 7 may result in the removal of a nuclear localization signal [46]. Our data show that modification of the C-terminal region of the ACT-like domain is sufficient to impair the bHLH protein’s ability to activate anthocyanin biosynthesis in all tissues, either through dimerization of the bHLH domain and consequent loss of RIF1 interaction or through abolishment of nuclear localization.

Interestingly, seed head color segregated 1:3 (green:purple) in an F_1_ population derived from the same parents. All F_1_ progeny displayed purple stolons and carried either the wild type MYC-bHLH allele or were heterozygous for the exon 7 (SNP_917_) mutation, as expected, because Meyer was heterozygous for the SNP_917_ mutation while PI 231146 was homozygous wild type. Our genetic analyses irrevocably showed that the loss of anthocyanin production in seed heads in 25% of the F_1_ progeny was caused by a second mutation in the same MYC-bHLH gene on chromosome 12. This mutation only affected anthocyanin production in seed heads. By definition, this mutation had to be heterozygous in PI 231146, and homozygous in both Meyer and the F_1_ parent of the F_2_ population (Fig. 6). All three lines had purple seed heads, and either homozygosity for this second mutation or heterozygosity at both the exon 7 mutation and the second mutation would result in green seed heads. Three SNPs were identified in the zoysiagrass MYC-bHLH gene that fulfilled these criteria (Table S7). SNP_281_ is located in the bHLH domain, SNP_716_ is not part of any conserved domain, and SNP_632_ is located in the N-terminal region of MYC-bHLH which physically interacts with MYB (Fig. S6) [47]. MYB transcription factors exhibit tissue-specific expression patterns that influence anthocyanin accumulation [48]. Because of the tissue-specific effect of the second mutation, we hypothesize that the G → T substitution at SNP_632_ eliminates binding of MYC-bHLH to a MYB transcription factor with seed head-specific expression, without substantially affecting interactions with MYB factors involved in the regulation of anthocyanin biosynthesis in stolons. A possible candidate is MYBc, encoded by *Zjn_sc00010.1.g01550.1.am.mk*, a close paralog to maize C1 (Fig. S7). C1 interacts with the MYC-bHLH transcription factor R and regulates anthocyanin production in the endosperm aleurone layer [49]. RT-PCR showed that *Zjn_sc00010.1.g01550.1.am.mk* is expressed in early seed heads but not in stolons or leaves (Fig. S9). In contrast, the *Z. japonica* ortholog to *C1*, *Zjn_sc00049.1.g03240.1.am.mk* (MYBa), lacks expression in both tissues, while its homoeolog *Zjn_sc00004.1.g00810.1.sm.mk* (MYBb) is expressed in both stolons and seed heads. Interestingly, our Y2H assays indicate that the MYBa and MYBb proteins derived from sc00049 and sc00004, respectively, interact much more strongly with both MYC-bHLH variants than MYBc (Fig. 5). There appears to be a slight reduction in the interaction of MYBa and MYBb with MYC-bHLH_Ser_ compared to MYC-bHLH_Ala_, but this reduction is much more pronounced for MYBc (Fig. 5). No effect of the MYBb and MYBc mutations in the interaction with either MYC-bHLH variant was observed. Further testing is needed to demonstrate that the Ala163Ser substitution in MYC-bHLH also results in a reduction and, potentially, loss of binding capability *in vivo*, and that the MYBb and MYBc factors tested are indeed involved in the production of anthocyanins in stolons and seed heads. However, the Y2H data confirm that the SNP_632_ mutation present in heterozygous condition in PI 231146 affects binding with MYB, and hence is likely causal, together with SNP_917,_ for the lack of anthocyanins observed in seed heads in the F_1_ population derived from Meyer and PI 231146 (Fig. 6).

**Figure 6.**
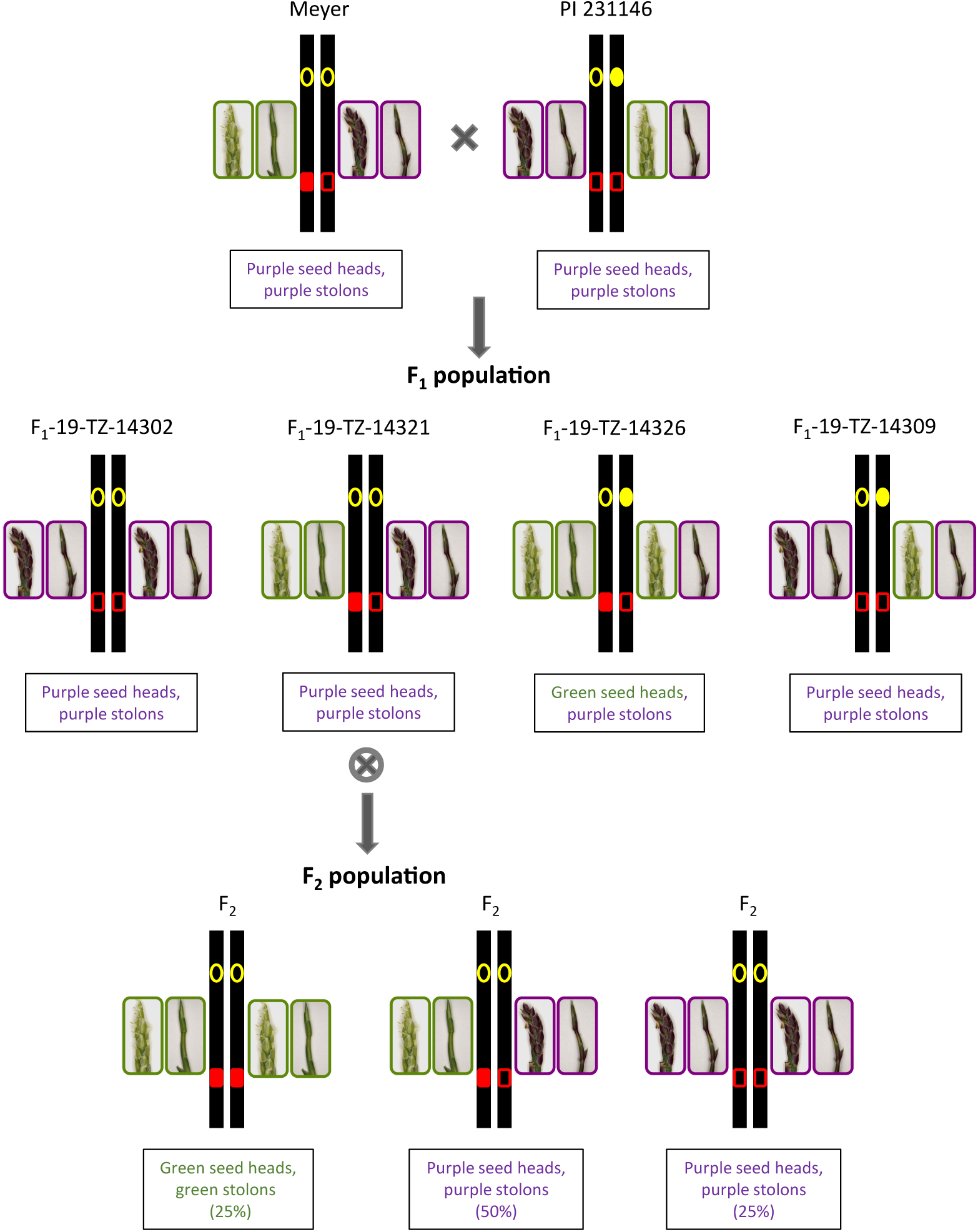
Schematic showing allelic composition and segregation of two MYC-bHLH mutations and the resulting anthocyanin phenotype associated with the haplotype in seed heads and stolons in the F_1_ (row two) and F_2_ (row three) populations. Yellow open and filled circles represent wild-type (G) and variant (T) at SNP_632_, respectively; red open and filled rectangles represent wild-type (G) and variant (A) at SNP_917_. In the F_1_ population, both variants segregate, while in the F_2_ population (selfed from F_1_ plant F_1_-19-TZ-14321), only SNP_917_ is segregating. SNP_917_ affects both seed head and stolon color, while SNP_632_ only influences seed head color. The IDs of the F_1_ plants provided are examples of progeny carrying the depicted allele combinations (Table S5).

## Conclusions

The *Z. japonica* x *Z. matrella* F_2_ maps will facilitate the identification of loci associated with desirable traits, specifically those that differentiate the two species. Although *Z. japonica* and *Z. matrella* are fully cross-compatible, several regions of extreme segregation distortion were observed. The complete elimination of *Z. matrella* alleles in the central region of chromosome 5 and the distal region of chromosome 7 indicates the presence of genetic drive factors. If the observations made in the Meyer x PI 231146 cross apply more widely to *Z. japonica* x *Z. matrella* interspecific crosses, introduction into *Z. japonica* of *Z. matrella* alleles linked to the drive loci will be extremely difficult. Reciprocal crosses need to be made to test involvement of the cytoplasm in the drive system.

The regulation of anthocyanin biosynthesis through the MBW complex has been well-studied with MYB factors frequently being identified as the primary regulators driving changes in this system [50–53]. However, in this study as well as a few others [19, 54, 55], mutations in MYC-bHLH transcription factors were identified as the underlying cause of anthocyanin loss in specific tissue types. What is unique in this study is that Meyer and PI 231146 were heterozygous for different mutations in MYC-bHLH that affected anthocyanin production in a tissue-dependent manner. The two mutations induced distinct structural, localization or interactive changes, leading to either complete loss of anthocyanin production (alteration of the ACT domain) or tissue-specific loss (in this case in seed heads) (likely caused by an amino acid substitution in the N-terminal domain). This highlights the dynamic nature of the MBW complex, wherein its composition, including the transcription factors involved, can vary depending on the tissue type. Knowing the mutations underlying anthocyanin production in zoysiagrass allows breeders to enhance turf’s aesthetic value, a highly desirable trait in the turfgrass industry, by achieving uniform color across leaves, stolons and seed heads.

## Materials and Methods

### Population development

In 2020, *Z. japonica* acc. Meyer was crossed with *Z. matrella* acc. PI 231146 at the University of Georgia (UGA), Tifton Campus. Subsequently, a validated F_1_ progeny, F_1_-19-TZ-14321, was selfed to generate ∼1000 F_2_ progenies. The F_2_ population was cultivated in 3x3x3 cm plastic pots in the greenhouse at UGA, Athens, and 530 random progenies were selected for genotyping and linkage map construction. A clonal duplicate of the population was planted in a maintenance nursery in Tifton, Georgia.

### DNA isolation, GBS library construction and sequencing

Healthy, fully opened leaves were collected from the two parents, F_1_-19-TZ-14321, and the randomly selected F_2_ progeny, and were flash-frozen in liquid nitrogen. The samples were stored at -80 °C until further processing. The frozen tissue was ground using a TissueLyser II bead mill (Qiagen), and the DNA was extracted using a modified CTAB method adapted from Doyle and Doyle [56]. DNA concentrations were measured with a Nanodrop, quality was assessed on a 1% agarose gel, and then each sample was diluted to 50 ng/µl. Subsequently, 200 ng of DNA was digested with the restriction enzymes *Pst*I-HF (NEB) and *Msp*I (NEB). Genotyping-by-sequencing (GBS) libraries were constructed as described in Qi et al. [23]. Each library was quantified using the dsDNA HS Assay Kit in a Qubit 2.0 Fluorometer (Invitrogen). Forty nanograms from each library, including for the F_1_ and the parents, were pooled. Approximately 200 libraries were combined into a single pool. Pools were subjected to Solid Phase Reversible Immobilization (SPRI) selection using Sera-Mag SpeedBeads to eliminate residual primers and small DNA fragments. Subsequently, pools were sequenced on an Illumina NextSeq2000 platform (paired-end 150 bp) at the UGA Georgia Genomics and Bioinformatics Core (GGBC).

### Sequence analysis and single nucleotide polymorphism (SNP) calling

The UGbS-Flex pipeline workflow and parameters described in Qi et al. [23] with a few modifications were used for sequence analysis and SNP calling. Briefly, following quality checks using FastQC v.0.11.8 [57], the pooled reads were demultiplexed using the ‘process_radtags’ function within the Stacks software [58]. The reads were then subjected to a series of processes, including removal of the barcodes, adapter sequences, restriction cut sites, and low-quality sequences using FASTX_trimmer from FASTX_toolkit v.0.0.14 (http://hannonlab.cshl.edu/fastx_toolkit). Alignment of the cleaned reads to the *Z. japonica* scaffold level reference genome (ZJN_r1.1) (https://zoysia.kazusa.or.jp/; [4]) was conducted using Bowtie2 with default parameters [59]. SNP calling was performed using the Genome Analysis Toolkit (GATK) [17] version 3.8 for the Unified Genotyper function (parameters –dcov 1000, −glm BOTH) and version 4.3 for the Haplotype Caller function (parameters-stand-call-conf 30 -ERC GVCF). The SNPs identified with both calling functions underwent filtering to retain only biallelic SNPs with a minor allele frequency > 5% and a quality depth value > 10. SNPs with a per sample read depth ≥ 8x were converted to mapping scores denoted as A (homozygous for the reference allele), B (homozygous for the alternate allele), H (heterozygous), D (A or H), and C (B or H) using an in-house Python script, SNP_Genotyper.v3.0.py, as described in Qi et al. [23]. The same script also consolidated SNPs within a GBS tag (1 kb region) [23]. The SNPs obtained with Unified Genotyper and Haplotype Caller were merged to generate a set of non-redundant SNPs.

### Genetic map construction

Only SNPs that were heterozygous in the F_1_ were retained for genetic mapping. Because of its outcrossing nature and because the alignment was done to the *Z. japonica* genome assembly, Meyer, the *Z. japonica* parent, was expected to be homozygous for the reference allele (A) or heterozygous while PI 231146, the *Z. matrella* parent, was expected to be homozygous for the alternate allele (B) or heterozygous at most of the loci. For markers that were B in Meyer or A in PI 231146 and segregated 1:2:1 in the F_2_ population, B scores were changed to A and *vice versa* in both the parents and progeny. Similarly, C scores were changed to D, and D to C. Further, because the linkage phase is unknown, markers that had H or missing data in either of the parents were duplicated, and A was changed to B, B to A, C to D and D to C in one copy. Markers with reversed scores were identified with the suffix ‘r’. Then, cosegregating markers were removed, retaining only a single representative marker, using an in-house python script ‘SNP_cosegregation.py’ (https://devoslab.franklinresearch.uga.edu/scripts-used-gbs-pipeline). Markers and progeny with more than 20% missing data were excluded. Finally, markers that exhibited segregation ratios deviating from Mendelian 1:2:1 ratios with p-values < 1x 10^-10^, as determined through chi-squared tests, were eliminated.

A combination of three genetic mapping procedures, MSTMap [60], SeSAM [61], and a version of MAPMAKER modified from Lander et al. [24] with a Python interface by Qi et al. [23], was employed for the construction of the linkage maps. MSTMap was employed to assign markers to linkage groups using a LOD score of 12 and a no-mapping threshold of 15 cM, resulting in a total of 32 linkage groups. Subsequently, duplicated linkage groups containing identical marker sets (following marker duplication and reversion) were excluded, leaving 22 unique linkage groups. Subsequently, the autoMap function (parameters - genotypingErrorDetection = TRUE, mappingFunction = “kosambi”) within SeSAM was employed to create a linkage map with robust marker orders. The population type was set to “F_2_”. The map was further refined by eliminating markers exhibiting more than 10 double recombination events across the 530 progenies. The final Kosambi distances between markers were calculated using MAPMAKER and associated Python scripts [23]. Marker orders were manually adjusted to agree with those in the *Z. japonica* ZJN_r1.1_pseudomol assembly only if reordering did not result in a net gain in recombination events.

### Comparing the genetic map with Zoysia and Eleusine coracana genome assemblies

BLASTN queries were executed using 1-kilobase regions centered on the mapped SNPs and extracted from the alignment of the GBS reads to the *Z. japonica* scaffold level assembly (ZJN_r1.1). These queries were applied against the *Z. japonica* cultivar Nagirizaki pseudomolecules (ZJN_r1.1_pseudomol; https://zoysia.kazusa.or.jp/), *Z. matrella* acc. Wakaba assembly (ZMW_r1.0; https://zoysia.kazusa.or.jp/), *Z. pacifica* acc. Zanpa assembly (ZPZ_r0.1; https://zoysia.kazusa.or.jp/) [4] and the high-quality KNE 796-S finger millet reference assembly (v1.1) [19]. For each query, the top hit (*Z. japonica* pseudomolecule and *Z. pacifica* assemblies) or top two hits (*Z. matrella* and finger millet assemblies) were recorded using an e-value threshold < 1 x 10^-5^. The top two hits for finger millet and *Z. matrella* had similar e-values, while there was generally only a single hit each for *Z. japonica* and *Z. pacifica*. Because the comparison of the genetic maps with the *Z. japonica* genome assembly showed that the distal regions of Chr02 and Chr07, and the central regions of Chr05 and Chr19 were missing from the genetic map, and that markers flanking the missing regions showed levels of segregation distortion close to or at the threshold used for removing highly distorted markers at the start of the mapping process (p-value < 1x 10^-10^), the scaffolds corresponding to the missing regions were identified, and markers were manually extracted. The maps for chromosomes 2, 5, 7, and 19 were then reconstructed using the mapping approach described earlier, without excluding markers based on segregation distortion.

In an attempt to differentiate the *Z. japonica* subgenomes, k-mer analysis was performed on the *Z. japonica* pseudomolecule assembly using k-mers of lengths 22–26, which were counted and grouped into sets with shared ancestry, independent of external genome comparisons, using the default parameters suggested for polyCRACKER [29].

### Phenotyping anthocyanin presence in stolons and seed heads

The presence of anthocyanins in seed heads was visually assessed in the greenhouse (UGA, Athens) in 137 flowering F_2_ progeny and in a nursery (UGA, Tifton) in 425 progenies. Because color intensity could be influenced by environmental factors, purple coloration was recorded as a binary trait (green = 1, purple = 2). Anthocyanin presence in stolons was determined in the greenhouse in 507 F_2_ progeny and similarly recorded. Pigmentation of seed heads was also scored in 153 F_1_ progeny derived from the same parents [16] and maintained in the Tifton nursery.

### Quantitative trait locus mapping

Quantitative trait locus (QTL) mapping for anthocyanin pigmentation was performed using R/qtl [62]. The analysis employed three marker covariates and utilized a Haley-Knott regression with a step size of 2.5 centimorgans (cM). Significance thresholds for the logarithm of odds (LOD) scores (p < 0.05) were determined by conducting 1000 permutations for each trait. The percentage variation explained (PVE) was calculated using the ‘makeqtl’ and ‘fitqtl’ functions [62].

### Whole genome shotgun sequencing of parents and F_1_ of the F_2_ population

DNA was extracted from Meyer, PI 231146 and F_1_-19-TZ-14321 as described for the GBS analyses. The DNA was digested with NEBNext dsDNA Fragmentase® enzyme (NEB) at 37 °C for 15 minutes, and whole genome libraries were made with the KAPA HyperPrep kit following the manufacturer’s guidelines but using half reactions. Library concentrations were measured with the dsDNA HS Assay Kit in a Qubit 2.0 Fluorometer (Invitrogen). Then, forty nanograms from each library were pooled and sequenced on an Illumina NextSeq2000 platform (paired-end 150 bp) at the GGBC.

### Candidate gene analysis

The QTL region was projected on the annotated finger millet KNE 796-S genome assembly, and genes in the QTL region were downloaded using Biomart in Phytozome (https://phytozome-next.jgi.doe.gov/info/Ecoracana_v1_1). Genes with a description or GO annotation in Phytozome containing the term ‘anthocyanin’ were further analyzed. Nucleotide variation in candidate genes was assessed from the whole genome sequencing data from the parents and F_1_ using SNPeff using default parameters [63]. SNPs of interest were narrowed down using the parameter view -i ‘INFO/ANN∼“HIGH” || INFO/ANN∼“MODERATE”’. Because comparative information showed that some *Z. japonica* acc. Nagirizaki gene models were likely incorrect, the intron/exon locations of all identified SNPs were manually verified using, if needed, adjusted gene models. Segregation in the F_2_ and F_1_ populations of two SNPs of interest was tested using cleaved amplified polymorphic sequence (CAPS) markers. For the *Dde*I CAPS marker (targeting SNP_917_), the primers used were E-E1F/ R (Table S9). For the *Bss*HII CAPS marker (targeting SNP_632_), the primers used were Pos2-F1/ R1 (Table S9). Intron retention caused by SNP_917_ was tested using primers E-E1F/ R. The annealing temperatures used are listed in Table S9.

### Phylogenetic analysis of select MYB transcription factors

The MYB transcription factors in finger millet (*Eleusine coracana* v1.1) most closely associated with maize C1 were used as BLASTP queries against the proteomes of *Z. japonica* acc. Nagirizaki and *Z. matrella* acc. Wakaba (zoysia.kazusa.or.jp). The top hits from these species were aligned using MUSCLE with default parameters in AliView [64]. Where necessary, gene structures from the *Zoysia* species were refined using the finger millet coding sequences, and the corresponding protein predictions were generated. Orthologs from rice (*Oryza sativa* v7.0) were also included in the alignment, alongside MYB transcription factors from *Arabidopsis* (AtTT2, AtPAP1, and AtMYB5) and maize (C1 and Pl). A neighbor-joining phylogenetic tree was constructed from the alignment using the MYB domain only in MEGA v11.0.13 [65]. The tree was built using the Jones-Taylor-Thornton (JTT) substitution model, uniform rates among sites, complete deletion for gaps and missing data, and 1000 bootstrap replicates.

## Semi-quantitative RT-PCR

To assess the expression of *Zjn_sc00049.1.g03240.1.am.mk* (MYBa), *Zjn_sc00004.1.g07010.1.sm.mk* (MYBb) and *Zjn_sc00010.1.g01550.am.mk* (MYBc; TT2), total RNA was extracted from leaf, stolon, and seed head tissues of select F_2_ samples using the TRIzol method. A total of 500 ng of RNA was converted to cDNA using Thermo Scientific’s RevertAid First Strand cDNA Synthesis Kit following the manufacturer’s protocol. The synthesized cDNA was diluted to 20 ng/μl, and 20 ng was used as a template for real-time (RT) semi-quantitative (sq) PCR. Each reaction consisted of 1X SsoAdvanced Universal SYBR Green Supermix (Bio-Rad) and 300 nM of gene-specific primers. The primers used were TT2_LP1/ RP3 for *Zjn_sc00010.1.g01550.am.mk,* MYB-manual-qF2/ qR1 for *Zjn_sc00004.1.g07010.1.sm.mk* and MYBhomeo-manual-qF1/ qR1 for *Zjn_sc00049.1.g03240.1.am.mk* (Table S9). Expression of a kinase gene (*Zjn_sc00058.1.g03630.1.am.mkhc*) using primers Kin-qF1/ qR2 (Table S9) was used as reference. PCR amplification was performed at 64 ^0^C for 28 cycles. The annealing temperatures used are listed in Table S9.

### Yeast two-hybrid assays

The Yeastmaker Transformation System 2 (Clontech) was used following the manufacturer’s instructions. Interaction tests were conducted in the yeast strain *Saccharomyces cerevisiae* Y2HGold (Clontech). To analyze interactions between MYC-bHLH and MYB, the first 756 bp of the coding region of the MYC-bHLH gene *Zjn_sc00004.1.g07010.1.sm.mk* (N-terminal 225 amino acids [66]) were synthesized (Twist Bioscience) and cloned into the pGBKT7 vector as bait. A second bHLH fragment was synthetized in which the SNP at position 487 (counting from the start codon; equivalent to SNP_632_) was changed from G to T. The full-length coding sequences of three MYB genes, *Zjn_sc00049.1.g03240.1.am.mk* (MYBa), *Zjn_sc00004.1.g07010.1.sm.mk* (MYBb; two variants differing by one SNP leading to an Asp76Gly substitution) and *Zjn_sc00010.1.g01550.1.am.mk* (MYBc; two variants differing by three SNPs leading to Gln203Arg, Arg256Gln and Ser279Ala substitutions) were cloned into the pP6 vector as prey. Because the annotated gene models as obtained from the Zoysia Genome Database (zoysia.kazusa.or.jp) were incorrect for several genes, the corrected coding sequences are provided in Fig. S10 and Fig. S11. Various bait and prey combinations were co-transformed into the Y2HGold yeast strain. Empty pGBKT7 and pP6 vectors were used as controls to test for auto-activation. Successfully co-transformed yeast cells were selected on synthetically defined (SD; Clontech) plates lacking tryptophan and leucine (SD/-Trp-Leu). Single colonies were inoculated into liquid SD/-Trp-Leu medium and cultured for 16–20 hours. Yeast cultures were adjusted to OD_600_ concentrations of 1.0, 0.1, and 0.01, and spotted onto minimal medium quadruple-dropout (QDO SD/-Leu-Trp-His-Ade) plates supplemented with either 50 µg/ml Aureobasidin A (Aba) (QDO/Aba), or 50 µg/ml Aba and 20 µg/ml X-α-Gal (QDO/Aba/XGal). Plates were incubated for five days at 30 ^0^C before being photographed.

## Acknowledgements

This research was supported by award #1915919 from the National Science Foundation Plant Genome Research Program to KMD, and award 2019-51181-30472 from the National Institute of Food and Agriculture (NIFA) – Speciality Crop Research Initiative (SCRI) to KMD, SM-L and BS. We thank Julia Ajello, Charlotte Greene, Gurjot Sidhu and Gowri Pillai for assistance with tissue collection and phenotyping.

## Data availability

Data supporting the findings of this work are presented within the main text, and as Supplementary Tables, Figures and Files. The whole-genome shotgun Illumina reads for *Z. japonica* acc. Meyer, *Z. matrella* acc. PI 231146 and F_1_-19-TZ-14321, and the GBS Illumina reads for Meyer, PI 231146 and the 530 F_2_ progeny used in the generation of the genetic map have been deposited to NCBI-SRA (Project PRJNA1235172).

## Author contributions

SP, JJS and KMD designed the experiments. SP conducted the GBS analyses, generated the genetic maps, conducted most of the phenotyping, carried out the QTL analyses and sequenced the parents. JZ conducted the Y2H experiments. EMB conducted the PCR analyses demonstrating the presence of two independent mutations in MYC-bHLH. BS generated the Meyer x PI 231146 F_1_ and F_2_ populations. JC validated the F_1_ progeny. XY and SM-L generated the GBS data for the F_1_ population. SK scored the F_1_ and F_2_ populations in the Tifton nursery for seed head color. KMD assisted with data interpretation. SP and KMD co-wrote the manuscript. All authors edited and approved the manuscript.

