## Supplementary material for "Two mutations in the same MYC-bHLH transcription factor cause segregation of purple coloration of stolons and seed heads in *Zoysia japonica* x *Zoysia matrella* F_2_ and F_1_ populations": Fig. S1- Fig. S11

**Supplemental Tables and Files:**

**Table S1** Linkage map of *Zoysia japonica* acc. Meyer x *Zoysia matrella* acc. PI 231146 F_2_ population, and comparative relationships

**Table S2** Scaffold orders based on the *Z. japonica* pseudomolecules (Tanaka et al., 2016) and the linkage map generated in this study, and synteny to finger millet

**Table S3** Percentage of colinear markers between zoysia chromosomes and syntenic finger millet A genome chromosomes

**Table S4** Stolon and seed head color of F_2_ progeny

**Table S5** Stolon and seed head color of F_1_ progeny and genotypic scores at two key mutations

**Table S6** Genes underlying the Chr12 QTL identified for anthocyanin variance in zoysiagrass, projected onto the finger millet KNE 796-S A genome.

**Table S7** Non-synonymous SNPs in *Zjn_sc00004.1.g07010.1.sm.mk* and their predicted effects using SNPeff

**Table S8** Genotypic scores for 29 random F_2_ progeny using a *Dde*I CAPS markers for Zjn_sc00004.1:2 965 917

**Table S9** Primers used in this study

**Supplemental File 1** Sub-genome phasing of *Z. japonica* acc. Nagirizaki using polyCRACKER with a k-mer size of 23.

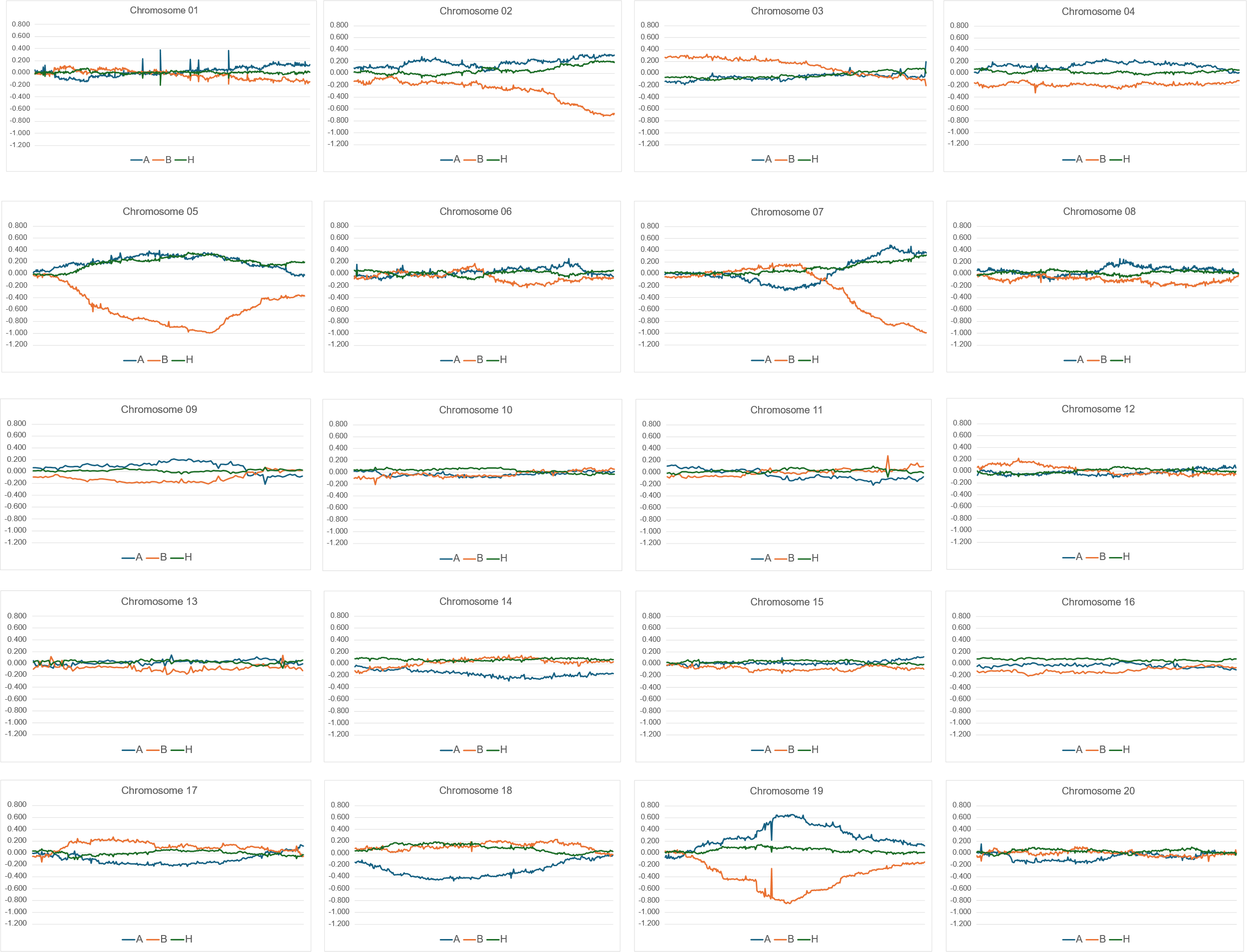

**Figure S1** Graphs showing segregation distortion in the *Z. japonica* acc. Meyer x *Z. matrella* acc. PI 231146 F_2_ population. For each allele, the ratio (Observed allele frequency – Expected allele frequency)/Expected allele frequency is shown along the chromosomes. Deviations from 0 indicate segregation distortion. Blue represents A alleles, orange represents B alleles, and green represents heterozygous genotypes.

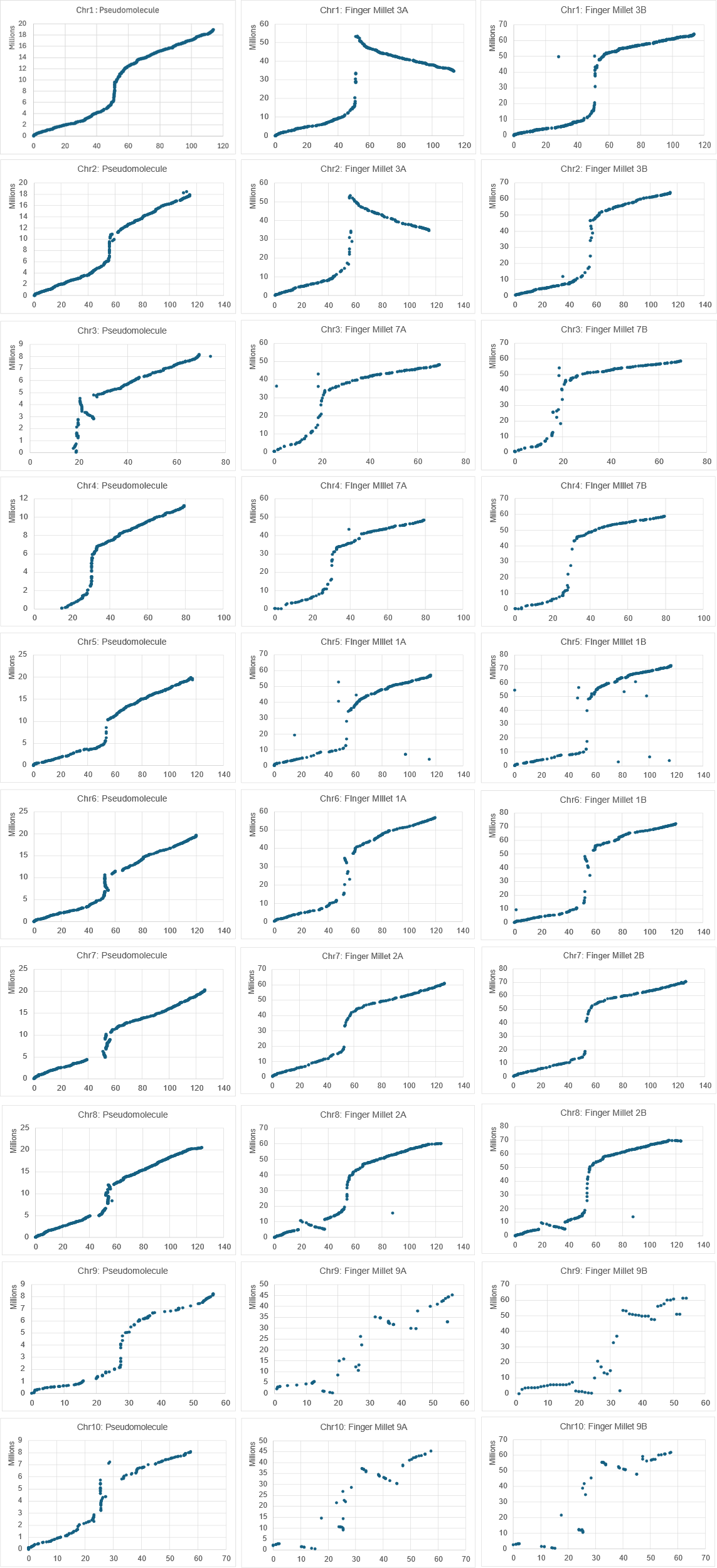

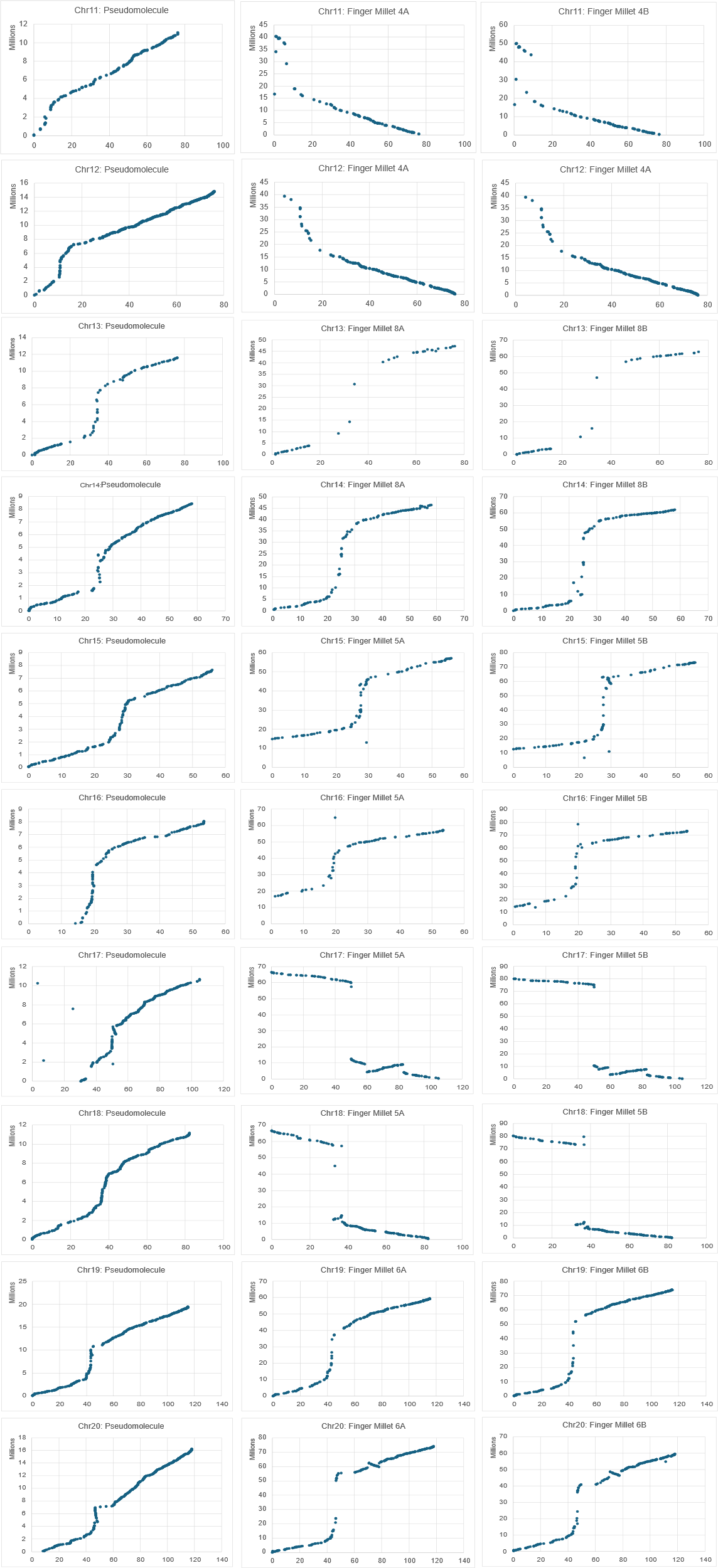

**Figure S2** Correlation between genetic distance (cM) (X-axis) and physical distance (bp) (Y-axis) across the *Zoysia* pseudomolecules (left panel), the *Eleusine coracana* accession KNE 796‑S subgenome A assembly (middle panel), and the *Eleusine coracana* accession KNE 796‑S subgenome B assembly (right panel)

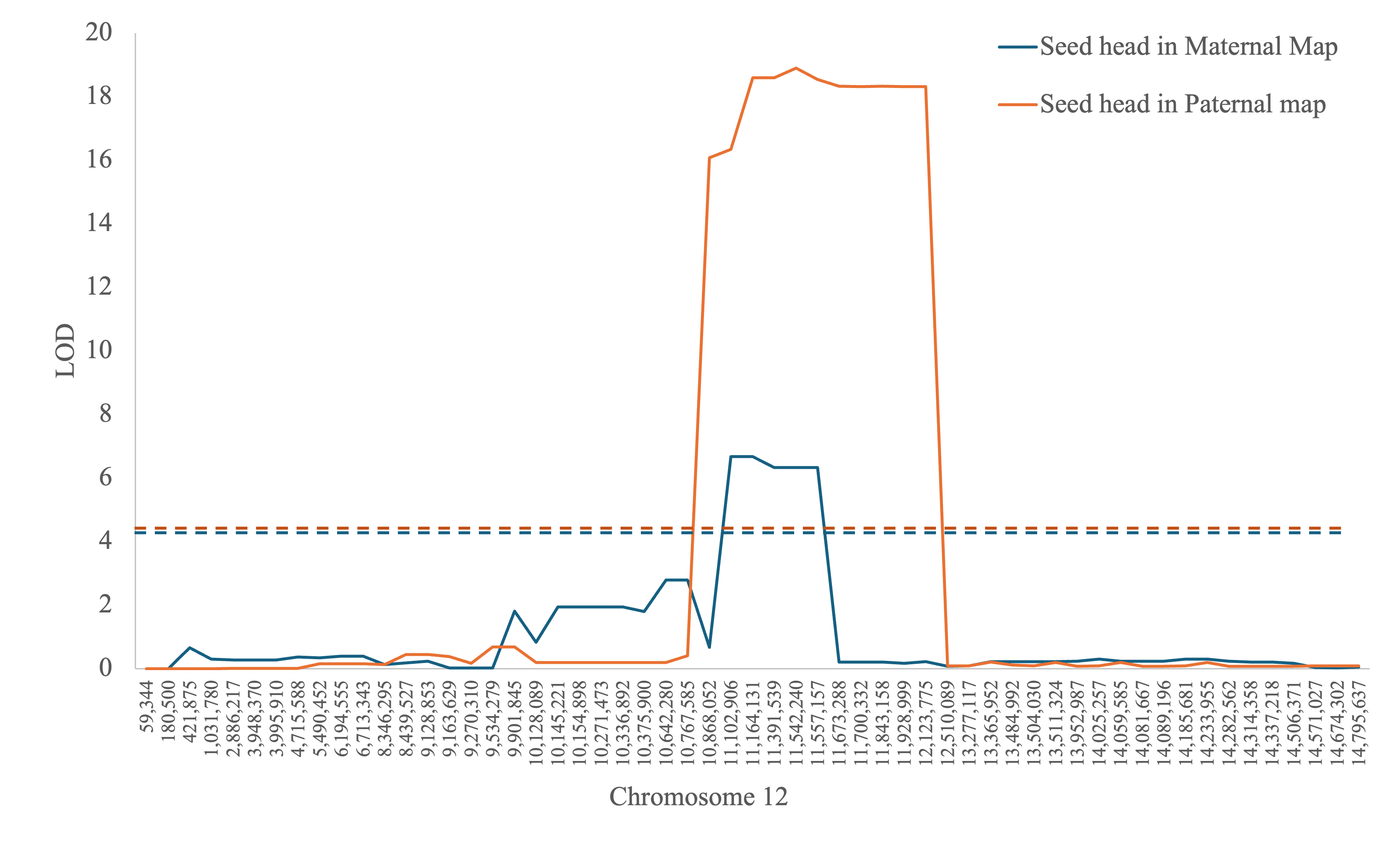
**Figure S3** Quantitative trait loci for anthocyanin pigmentation on chromosome 12 in the Maternal (Meyer) linkage map (blue) and Paternal (PI 231146) linkage map (orange). X-axis: Position on *Z. japonica* pseudomolecule (bp). Y-axis: LOD (logarithm of the odds)

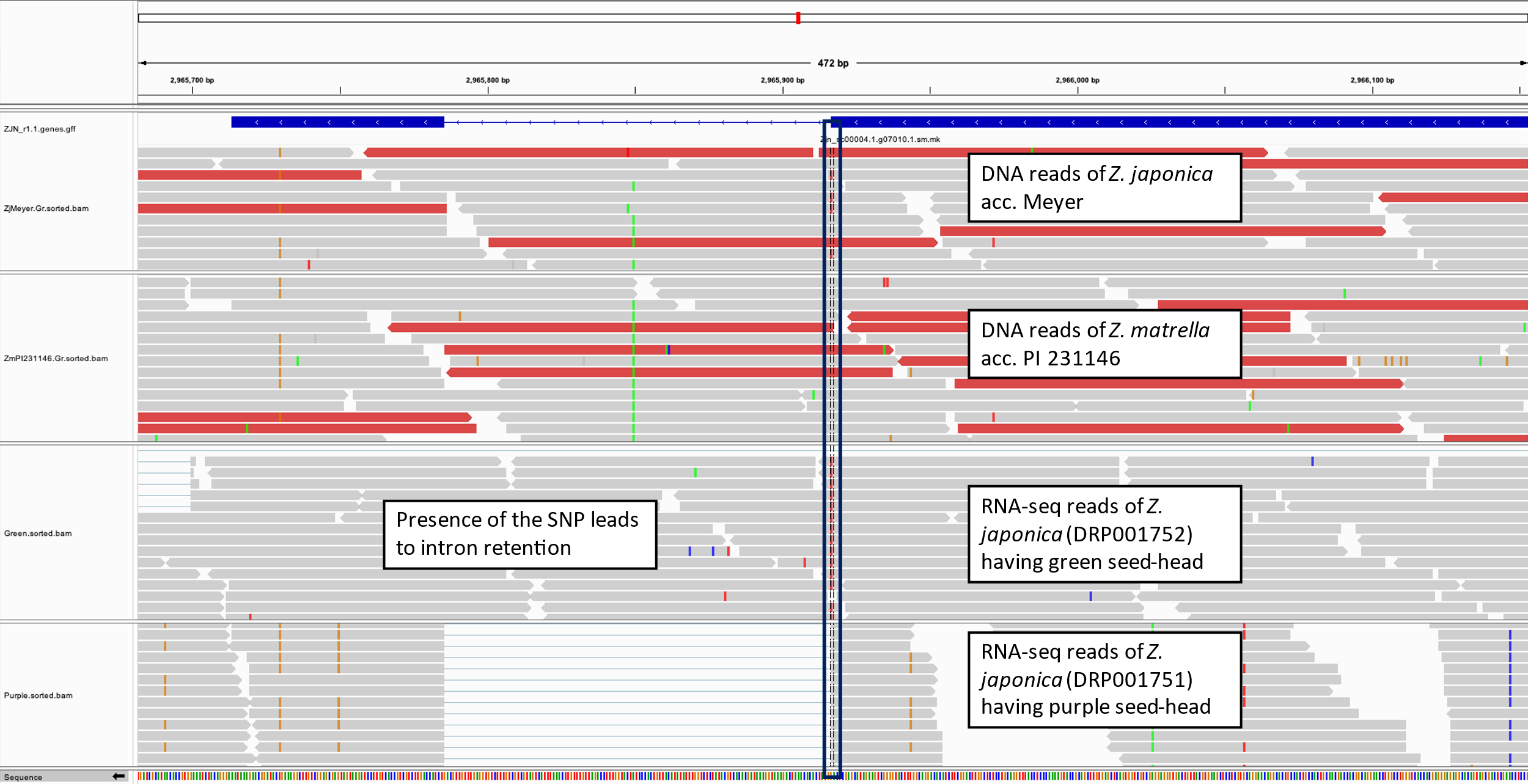
 **Figure S4** Integrative Genomics Viewer (IGV) snapshot of G → A SNP (black rectangle) in the MYC-bHLH encoding gene *Zjn_sc00004.1.g7010.1.am.mk*. The reference genome used in the alignment is the scaffold level assembly of *Z. japonica* cultivar Nagirizaki. The read sets (from the top) are whole genome Illumina reads of Meyer, whole genome Illumina reads of PI 231146, RNA-seq reads (SRA DRA001679) of a *Z. japonica* accession with green seed heads, and RNA-seq reads of a *Z. japonica* accession with purple seed heads (Ahn et al., 2015).

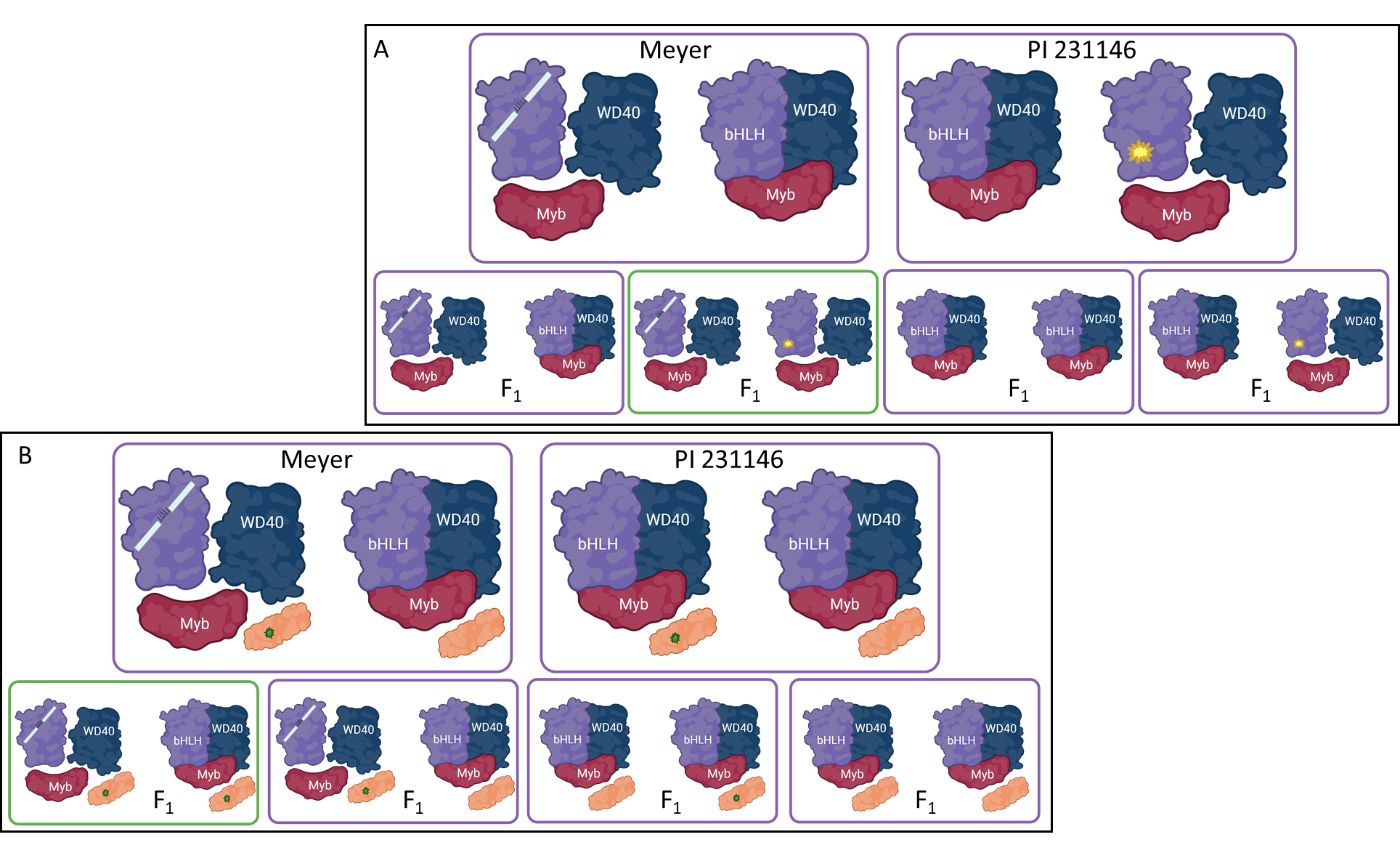
**Figure S5** Schematic showing expected segregation patterns in the F_1_ population when (A) Meyer and PI 231146 are heterozygous for two different mutations in the same *MYC-bHLH* gene (hypothesis 1) and (B) Meyer is heterozygous for a mutation in *MYC-bHLH* (indicated by a white bar representing an intron; mutation inactivates both stolon and seed head anthocyanins when homozygous) as well as a mutation in a linked gene (gene 2; in orange; mutation represented by a green star; mutation inactivates only seed head anthocyanins when homozygous) and PI 231146 is heterozygous for the same mutations in gene 2 (hypothesis 2). For hypothesis 2, the two mutations could be on the same chromosome in Meyer (as shown here), in which case the F_1_ that gave rise to the F_2_ population should have the same variant configuration as Meyer for stolon and seed head color to cosegregate, or across the two homologous chromosomes (not shown), in which case the F_1_ that gave rise to the F_2_ population should be heterozygous for the intron-retention mutation and homozygous wild-type for the second mutation. The white bar in bHLH indicates the mutation leading to intron retention. The yellow and green stars indicate SNP variants. The color of the box (purple or green) indicates the expected seed head color for each configuration. Figures were generated using Biorender (Biorender.com)

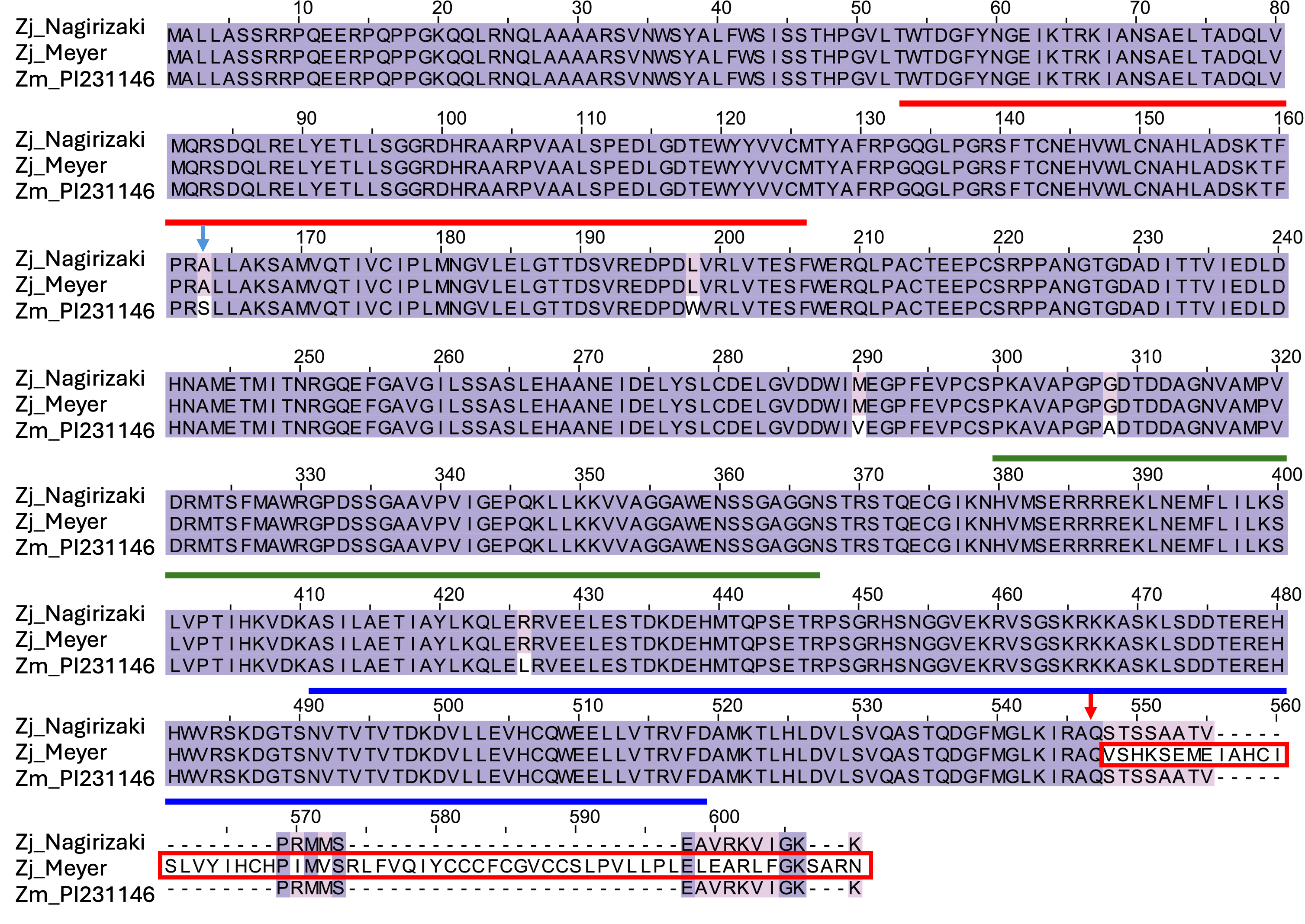

**Figure S6** Protein-level multiple sequence alignment of MYC-bHLH encoded by *Zjn_sc00004.1.g07010.1.sm.mk.* From top to bottom:  The wild type protein (from *Z. japonica* acc. Nagirizaki genome assembly), Meyer variant protein (position of intron retention and alteration of the C-terminal region of the protein (red box) caused by the SNP917 variant is indicated with red arrow), and PI 231146 variant protein (five amino acid substitutions caused by mutations at SNP_632_, SNP_045_, SNP_770_, SNP_716_ and SNP_281_; see Table S7 for SNP designations). SNP_632_, which may affect MYB binding, is indicated with a blue arrow. The N-terminal domain is indicated with a red horizontal bar, the bHLH domain with a green horizontal bar and the ACT domain with a blue horizontal bar.

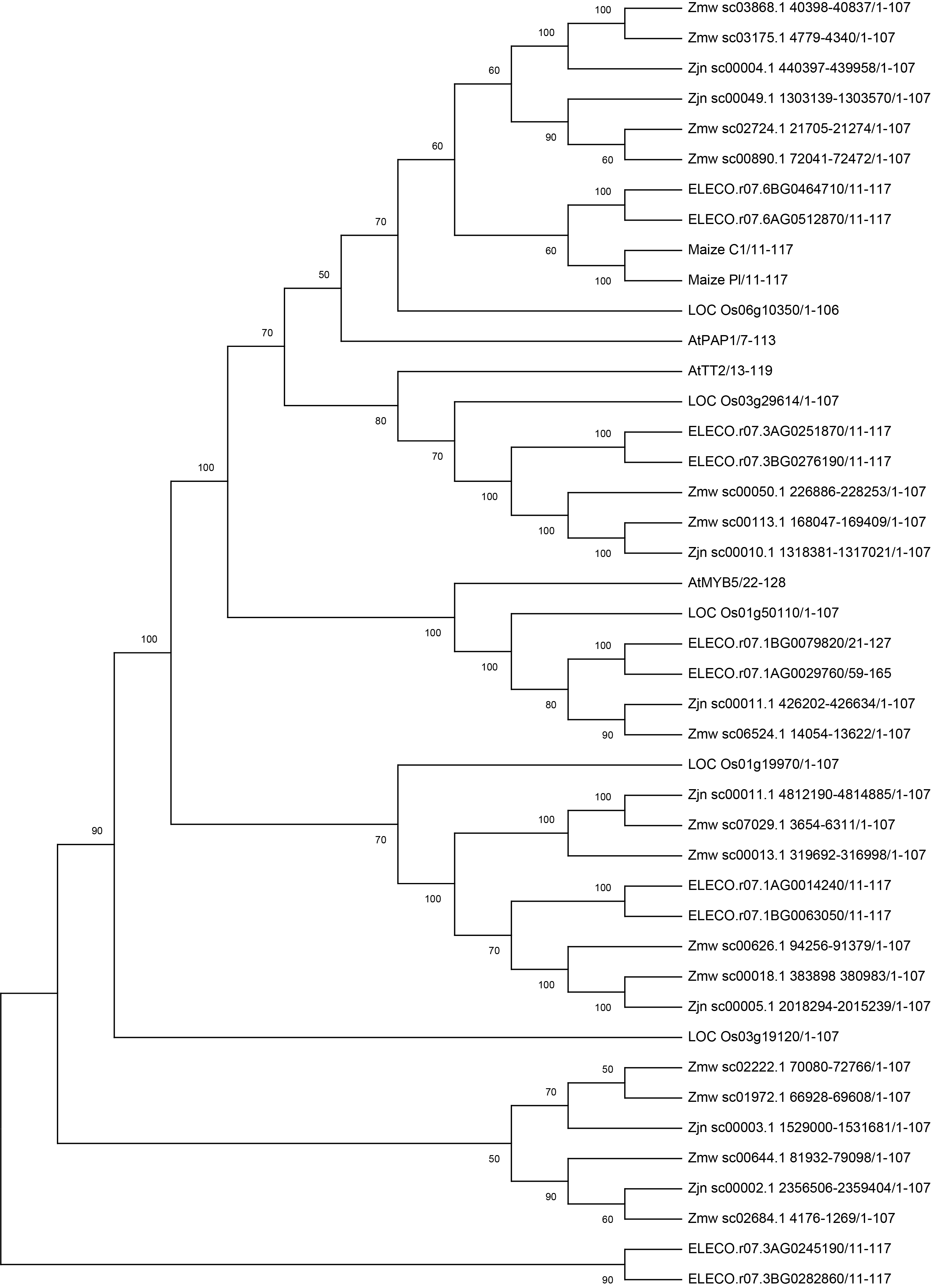
**Figure S7** Phylogenetic relationship of MYB proteins potentially involved in the anthocyanin pathway. Neighbor Joining tree showing the relationship between select MYB family members from *Arabidopsis* (At), finger millet (ELECO), rice (Nipponbare; LOC), *Zoysia matrella* (Wakaba; Zmw) and *Zoysia japonica* (Nagirizaki; Zjn). Support for the topology of the tree is provided by bootstrap values (1000 repetitions) at each node.

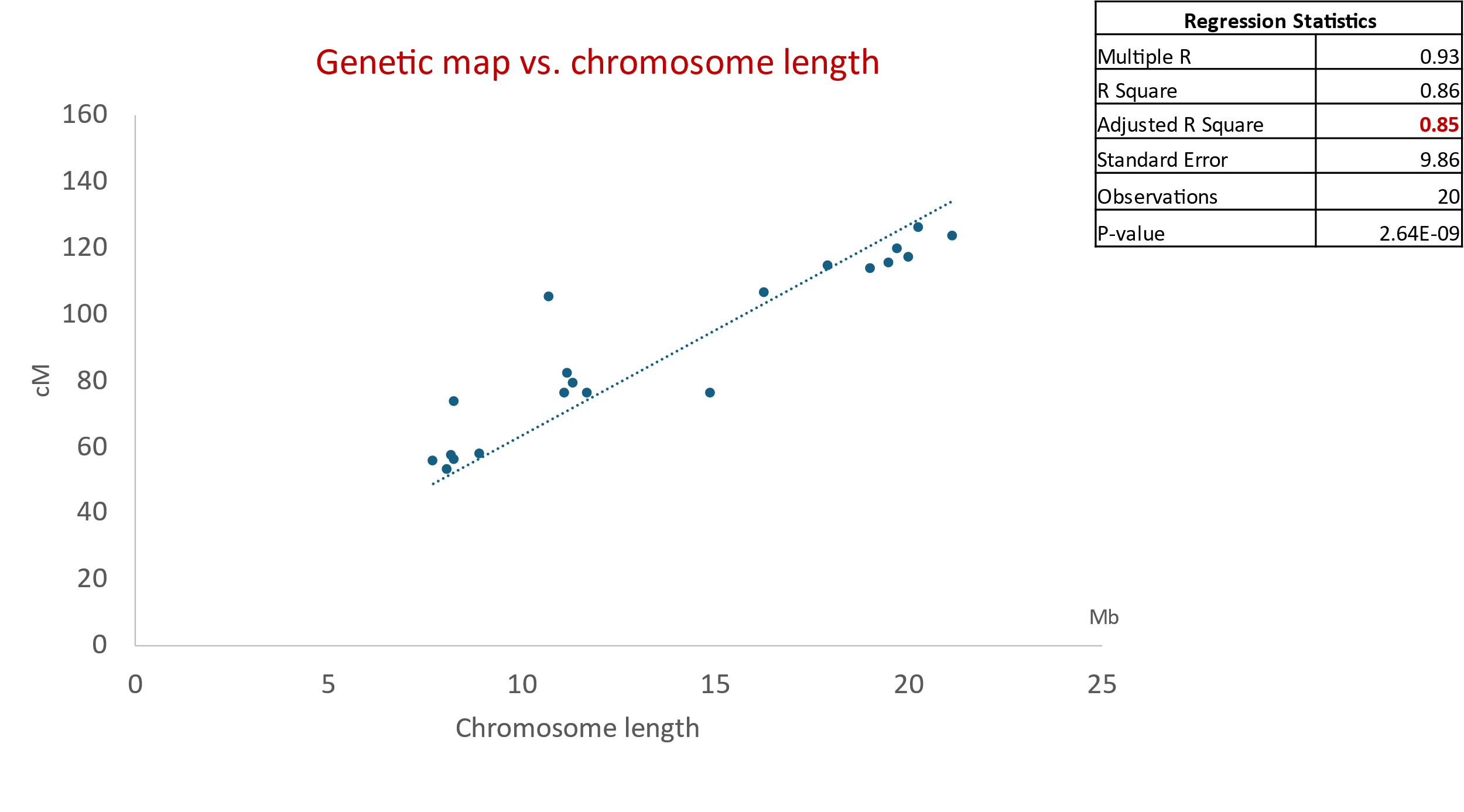
 **Figure S8** Correlation between the length of each linkage group in the F_2_ genetic map (in cM) (Y-axis) *vs.* the physical length (in Mb) of the corresponding *Z. japonica* chromosome (X-axis)

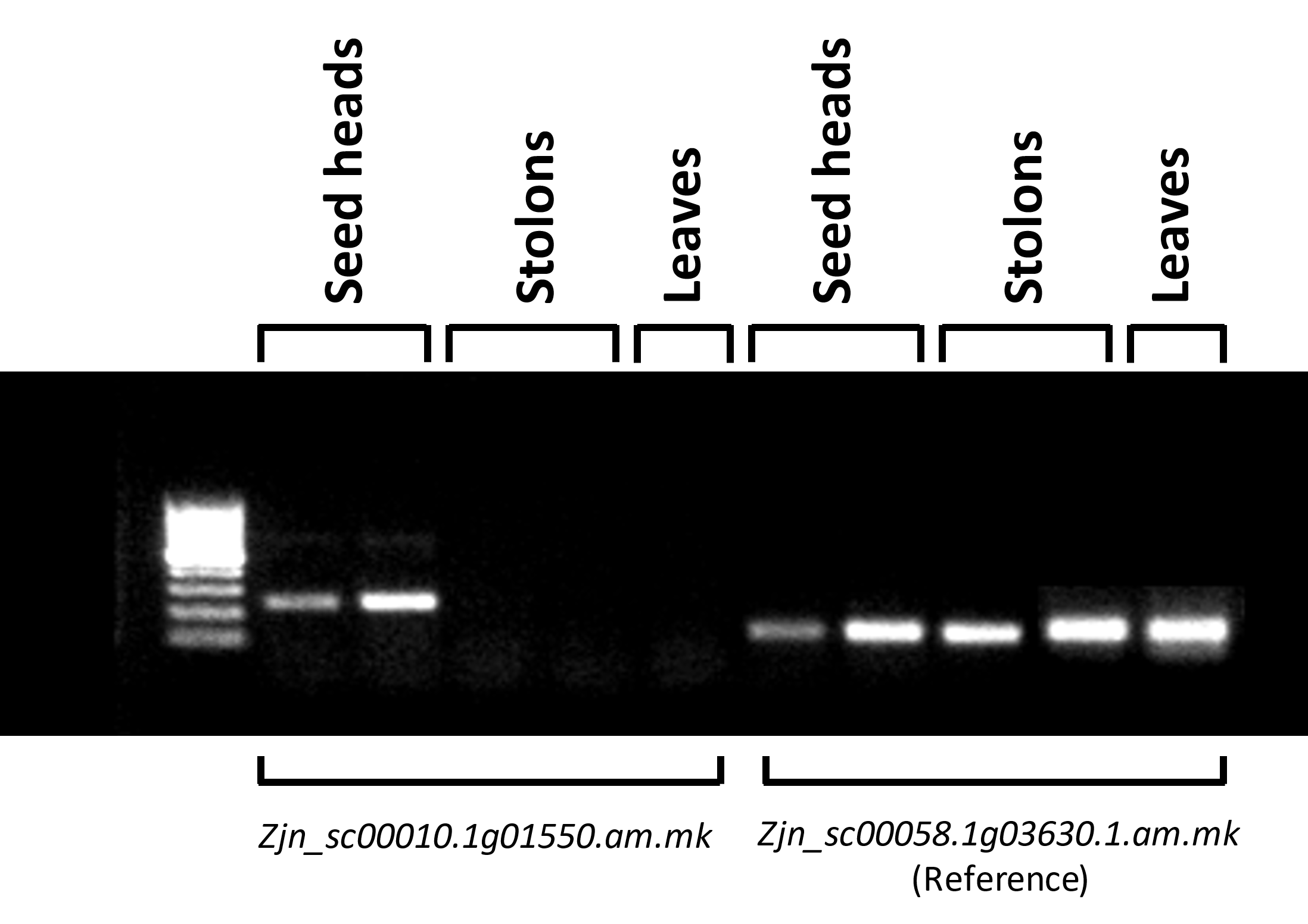

**Figure S9** Semi-quantitative-RT-PCR showing expression of *Zjn_sc00010.1.g01550.1.am.mk* (lanes 2-6) and the reference gene *Zjn_sc00058.1.g03630.1.am.mkhc* (lanes 7-11) in different tissues. Lane 1: 100 bp marker; Lanes 2 and 7: seed head sample 1; Lanes 3 and 8: seed head sample 2; Lanes 4 and 9: stolon sample 1; Lanes 5 and 10: stolon sample 2; Lanes 6 and 11: leaves. Primers used are listed in Table S9.

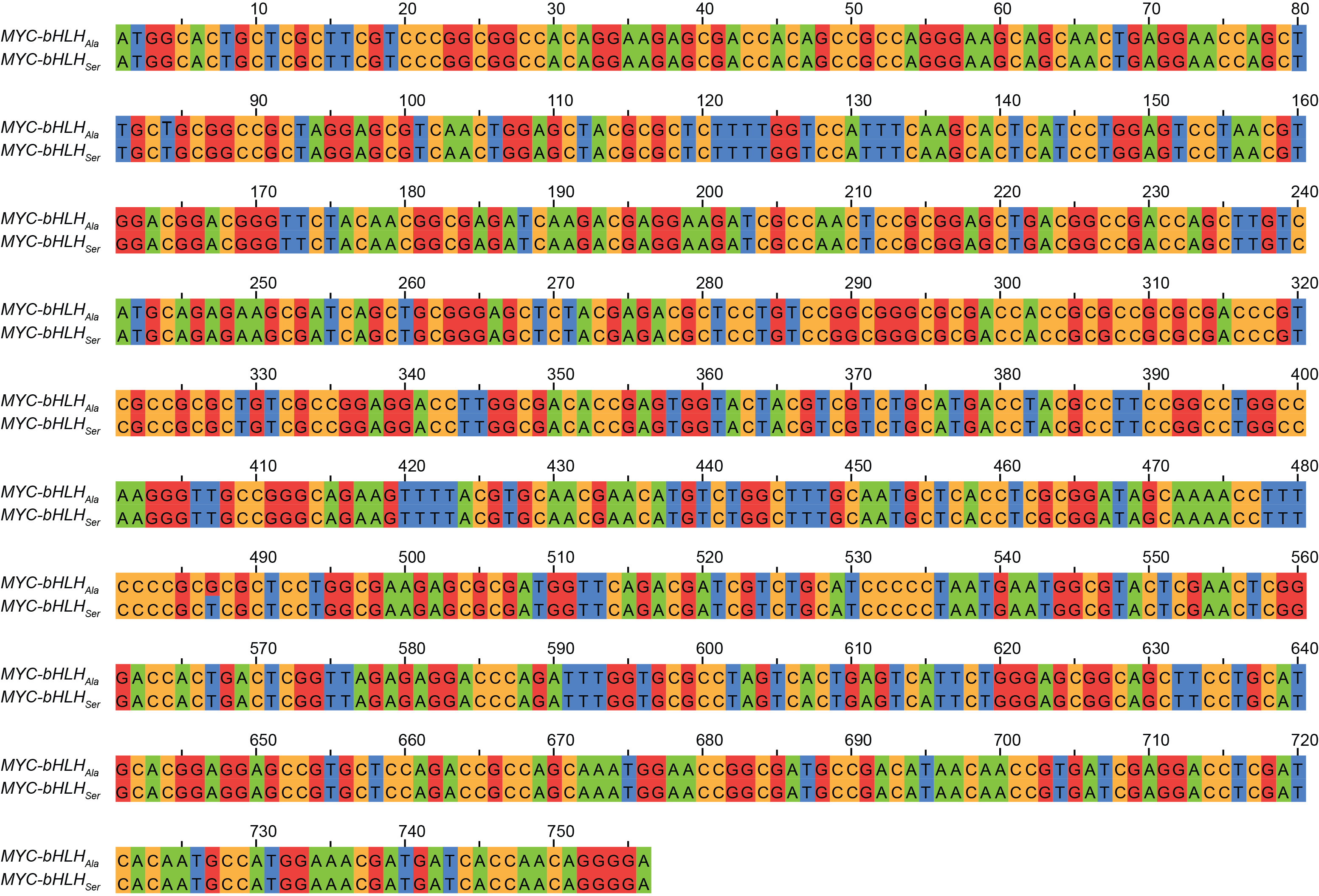

**Figure S10** MYC-bHLH_Ala_ and MYC-bHLH_Ser_ cDNA fragments used as bait in Y2H experiment

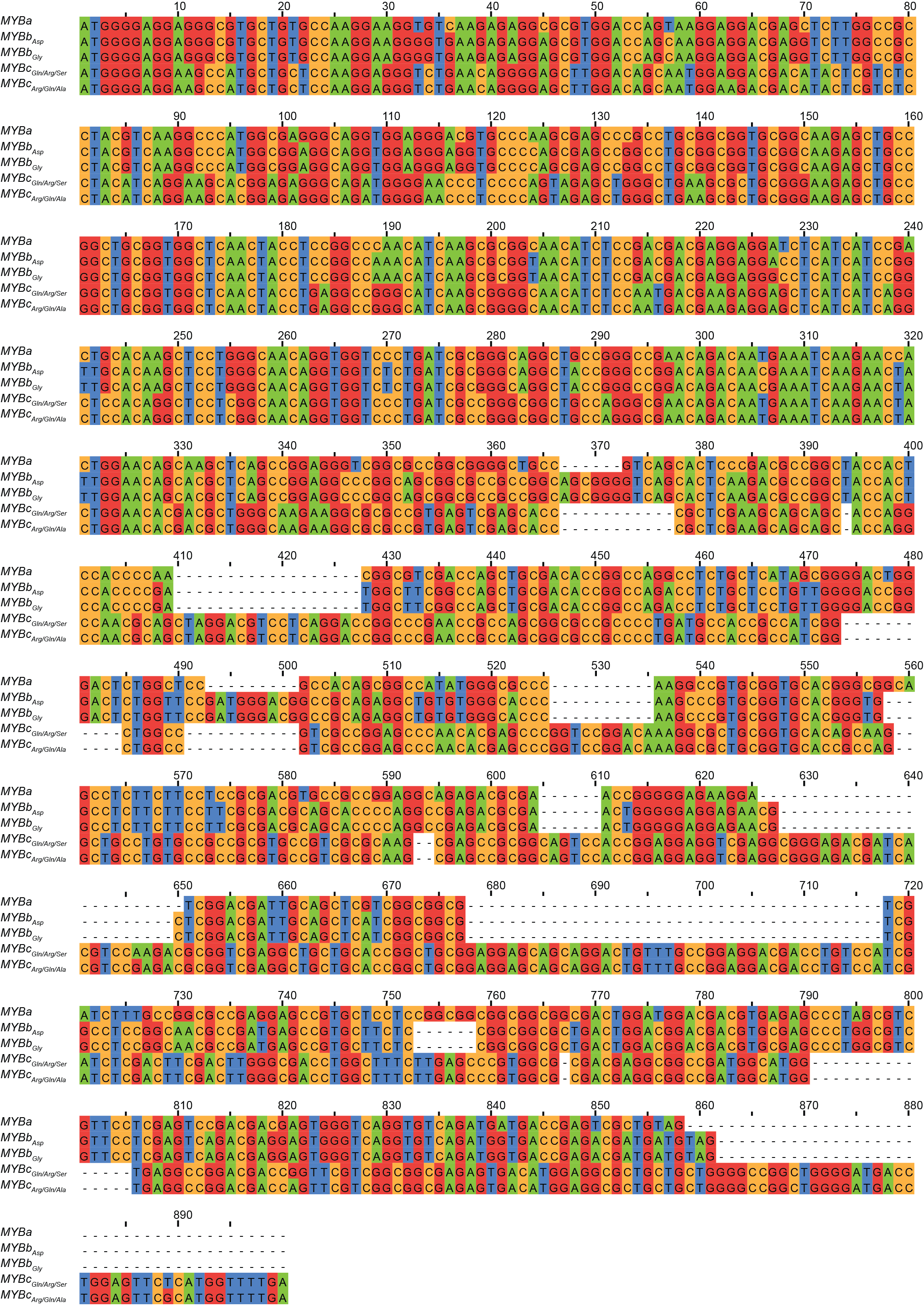

**Figure S11** MYBa, MYBb_Asp_, MYBb_Gly_, MYBc_Gln/Arg/Ser_ and MYBc_Arg/Gln/Ala_ cDNA sequences used as prey in Y2H experiment
